# Genomic context tunes sequence-encoded RNA processing

**DOI:** 10.64898/2026.10.29.762830

**Authors:** Mathias Nielsen, Artyom Tyukaev, Lidia Garkul, Igor Ulitsky

## Abstract

Sequence and genomic position jointly determine RNA processing, but at native loci the two are difficult to separate. Here, we uncoupled them by integrating RNA-processing elements into endogenous transcription units at tens of thousands of mapped positions and different elements at shared positions. Genomic position dominated RNA production levels but differentially influenced RNA processing events. A cryptic polyadenylation site, used natively only upon U1 inhibition, was rarely used ectopically. In contrast, introns partially retained at their native loci remained so across thousands of ectopic positions, with a substantial fraction of transcripts reaching the cytoplasm unspliced. SON depletion reduced splicing of two such introns, preferentially near speckles. Chromatin state, GC content, and speckle proximity tuned this behavior without overriding it, and in opposite directions: speckle proximity improved the splicing of one intron and reduced that of another. Specialized introns are therefore portable modules carrying their own processing information.

## Introduction

RNA processing is governed by an interplay between cis-regulatory sequence motifs and the local genomic environment in which those sequences are transcribed. Splicing, cleavage and polyadenylation (CPA) are specified by local sequence elements, but occur co-transcriptionally and are influenced by chromatin and nuclear contexts (Luco et al. 2010; Naftelberg et al. 2015). Transcriptional activity, RNA Pol II elongation rate, local sequence composition, subnuclear positioning, and chromatin state have all been associated with the regulation of RNA processing and alternative splicing (de la Mata et al. 2003; Bentley 2014; Moreau et al. 2025). Dissecting the contributions of each of these is particularly difficult because genomic features are highly correlated: gene-dense regions, for example, tend to be GC-rich, contain shorter introns, be enriched for highly and broadly expressed housekeeping genes and occupy more interior nuclear regions (Tammer et al. 2022). Therefore, a persistent challenge has been to separate the intrinsic biology of RNA-processing elements from the nuclear environment in which they normally occur. Beyond the effects of the local genomic neighborhood on RNA processing, the spatial positioning of the gene in the nucleus plays an important role. In particular, biomolecular condensates are emerging as important regulators of transcription and RNA processing (Pei et al. 2025). Nuclear speckles are enriched in splicing factors and are organized around the scaffold proteins SON and SRRM2 (Faber et al. 2022; Ilik et al. 2020), and proximity to nuclear speckles has been associated with increased splicing efficiency (Ding and Elowitz 2019; Bhat et al. 2024).

The functional importance of nuclear speckles is not fully clear, but they appear to be related to the phenomenon of intron retention. A class of partially retained introns requires nuclear speckle proximity and functionality for accurate processing has been identified (J. Wu et al. 2024; Małszycki et al. 2026; Wen et al. 2026; Barutcu et al. 2022). These introns share defining features: they are GC-rich, shorter than average, and enriched in genes involved in RNA processing and cell-cycle regulation, with retention of some varying across the cell cycle (Dominguez et al. 2016). Many overlap with the previously defined class of detained introns, suggesting that they may be spliced in response to environmental signals (Sung et al. 2023; Boutz et al. 2015; Barutcu et al. 2022), and are enriched among extensively conserved intronic regions (Farhangmehr et al. 2026). Their retention not only alters splice-site choice but also determines the fate of the host transcript, targeting it for nonsense-mediated decay (NMD) (Kurosaki et al. 2019) or restricting its export (Braunschweig et al. 2014; Boutz et al. 2015), although some intron-retaining transcripts reach the cytoplasm (Monteuuis et al. 2019). Whether the characteristic processing of these introns is encoded locally within the intron-containing sequence, or instead depends on the native host gene, chromatin environment or nuclear position, remains unresolved.

Premature cleavage and polyadenylation (PCPA) is another example of an RNA-processing event that may be jointly controlled by sequence and genomic context (Mitschka and Mayr 2022; Geisberg et al. 2024). Cleavage at intronic PCPA sites terminates transcription before canonical full-length transcripts can be produced. Although PCPA sites are widespread within transcription units, only a subset is used, indicating that the presence of a polyadenylation signal does not guarantee transcription termination. The U1 snRNP plays a central role in suppressing PCPA and is required for full-length transcription at thousands of genes (Venters et al. 2019; Kaida et al. 2010; Berg et al. 2012) by inhibiting access of CPA machinery to PCPA sites during transcription and promoting Pol II elongation (Mimoso and Adelman 2023; Oh et al. 2017). Still, the extent to which the activity of an individual PCPA site is encoded by its local sequence or determined by genomic context remains unclear.

Massively Parallel Reporter Assays (MPRAs) have emerged as a powerful approach for studying regulatory sequences at scale, but many conventional MPRAs use episomal plasmid reporters or integrate reporters at a single site (Mikl et al. 2019). A relevant approach for studying genomic context effects is Thousands of Reporters Integrated in Parallel (TRIP) (Akhtar et al. 2013), which enables high-throughput analysis of a single reporter across thousands of distinct genomic locations through piggyBac (PB) mediated random integration. PB integration of splicing cassettes has also formed the basis for gene-trap capture of endogenous transcription units through integration-dependent splicing (S. Wu et al. 2007; W. Wang et al. 2009; Pettitt et al. 2013); more recently, random integration of functional tags flanked by a splice donor has been used for large-scale tagging of the proteome (Nemčko et al. 2024), highlighting the advantage of coupling exogenous sequences directly to native local transcriptional units for functional studies.

Here, we combined these principles to develop promoterless single-exon and intron-containing TRIP reporters, whose expression relies on integration into existing transcription units, and used them to deconvolute the sequence-autonomous effects on RNA processing from those of the genomic context at tens of thousands of defined genomic positions. Genomic context strongly determined transcriptional output and quantitatively modulated RNA processing in a sequence-specific manner. A PCPA site that is natively U1-suppressed was essentially inert when inserted in thousands of genomic locations, whereas speckle-associated partially spliced introns preserved their characteristic processing across genomic locations. Intron identity shaped the subcellular distribution of the host transcript and conferred differential sensitivity to SON depletion and nuclear speckle proximity. Intron-containing sequences therefore carry portable regulatory information that determines processing efficiency, but this information has a variable impact that is affected by the genomic neighborhood and the nuclear environment.

## Results

### Promoterless TRIP reporters enable systematic measurement of position-dependent RNA processing

Reporter assays are widely used to dissect how sequence features regulate RNA processing, but transfected plasmids do not capture the influence of the chromatin and the surrounding genomic environment. To systematically examine how sequence and chromatin context jointly shape RNA processing, we adapted the Thousands of Reporters Integrated in Parallel (TRIP) method (Akhtar et al. 2013) with single-exon reporters integrated across thousands of genomic locations.

We first designed two piggyBac (PB) transposon system single-exon reporters to investigate the influence of genomic context on RNA processing. Each reporter contains a fixed 832 bp sequence that includes 64 bases from the end of the first intron and the full second exon of β-globin, where we inserted a unique 18 bp barcode and an additional exogenous sequence, and one of two 3’ intronic regions, all flanked by PB inverted terminal repeats (ITRs) (Fig. 1A). The variable intronic region consisted of either a 61 bp (‘Regular’ reporter) or a 1370 bp region derived from the first intron of *GNAI1* containing a premature cleavage and polyadenylation (‘PCPA’ reporter). In its endogenous context, this PCPA signal is suppressed by U1 snRNP; however, it becomes de-repressed upon the inhibition of U1 snRNP recognition via U1 antisense morpholino oligonucleotides (AMOs) (Fig. S1A). The plasmids were transfected into K562 cells, along with a plasmid expressing the PB transposase to facilitate random genomic integration of the reporter sequence.

**Figure 1.**
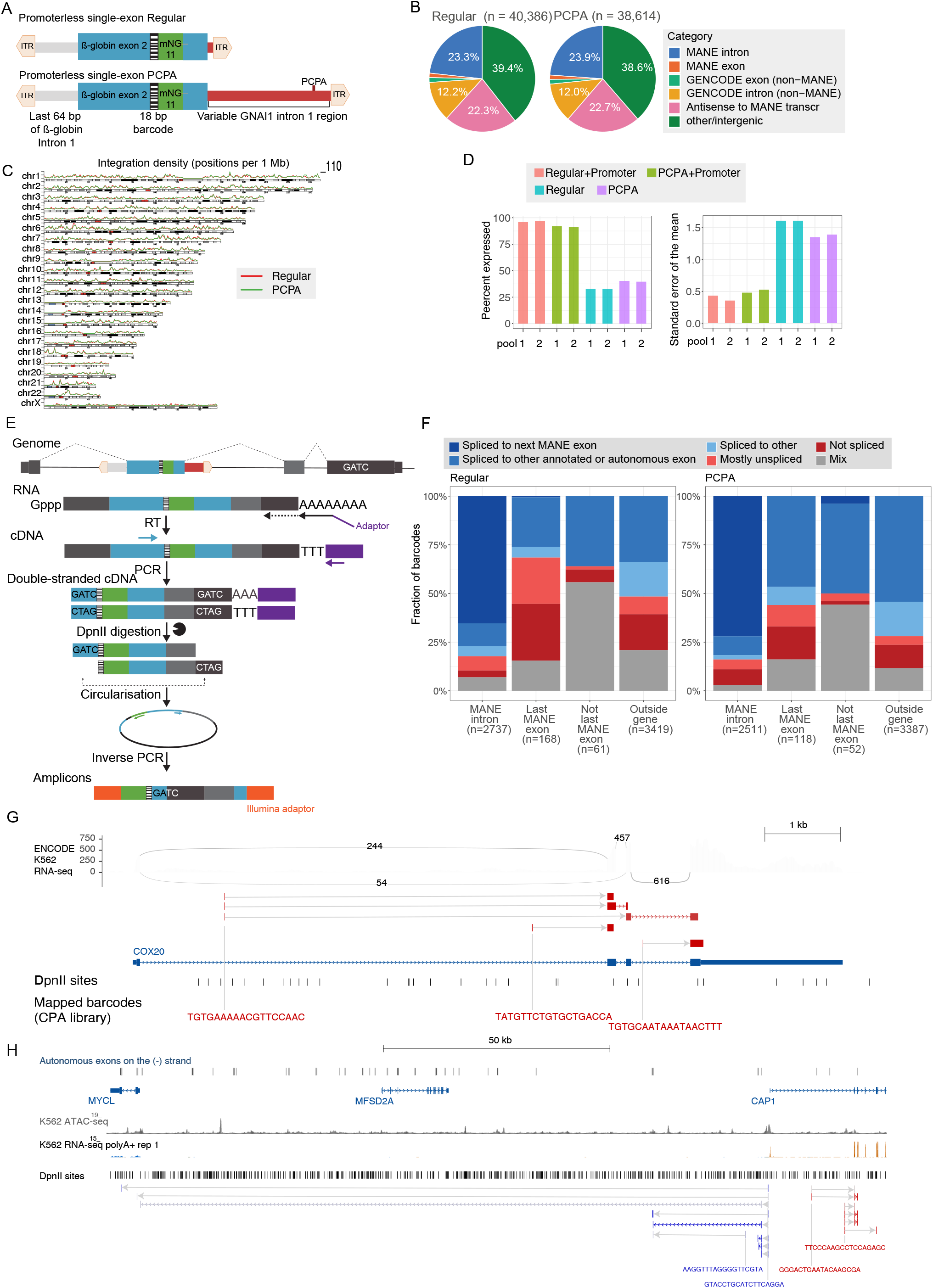
TRIP method for measuring genome-wide effects of single exon expression. **(A)** Outline of the Regular and PCPA reporters’ design. **(B)** Fraction of barcodes in each library that map to the indicated genomic region. **(C)** Density of integrations along human chromosomes. **(D)** Left: fraction of barcodes in the indicated library and pool that were expressed; Right: Standard error of the mean of log10-transformed expression values. **(E)** Overview of the method for mapping RNA sequences downstream of the integrated reporter, exemplified by an intronic integration. Gray exons indicate host transcript exons, with one of them containing, by chance, a GATC tetramer sequence. RNA was reverse transcribed with an oligo-dT primer containing an adaptor sequence (purple); a reporter-specific primer was then used to make double-stranded cDNA. The ds-cDNA was then digested with the 4-bp (GATC) cutter DpnII, which cuts upstream of the barcode in the reporter and at endogenous recognition sites expected on average every ∼250 nt. Digested ds-cDNA was then ligated at low concentration to facilitate intramolecular ligation, and sequence amplicons were made through inverse PCRs, resulting in a final amplicon that allows the mapping of the reporter barcode and the next downstream host exonic sequence (gray). **(F)** Distribution of reporter-host connectivity patterns observed for individual barcodes for which at least five informative reads could be obtained. **(G)** Barcodes mapped to the *COX20* gene body with informative reads from the reporter-host connectivity libraries. Gray arrows connect barcode location to the beginning of the RNA mapping reads. Regular RNA-seq from K562 cells from the ENCODE project is shown on top, with numbers on the arcs representing the numbers of splice-mapping reads. **(H)** Examples of barcodes in the gene desert between CAP1 and MYCL that are spliced to autonomous exons. Barcodes and RNA reads mapping to the plus strand are in red, and those mapping to the minus strand are in blue, with element intensity correlated with the number of reads supporting that splicing event. Above, ATAC-seq and stranded RNA-seq (red=(+) strand; blue=(-) strand) from the ENCODE project for K562 cells are shown.

The transfected cells were cultivated for 7 days, and two TRIP pools from each construct were established and subcultured for an additional ∼2 weeks as in the original study ((Akhtar et al. 2013). Genomic DNA was extracted, and insertions were mapped through inverse PCR followed by high-throughput sequencing. We mapped 40,386 insertions from the Regular library and 38,614 from the PCPA library, which were assigned to 39,816 and 38,074 unique genomic positions, respectively, with median distances of ∼32 kb between integrations (Table S1 and S2). Among the initially mapped positions, more than 90% were within the known PB tetranucleotide TTAA, and only uniquely mapped barcodes that were also identified in amplicon libraries prepared from genomic DNA (see below) were analyzed further. We randomly selected four insertions and validated their integration positions by PCR (Fig. S1B). Insertions were identified across the genome, with a preference for open chromatin regions and expressed genes, as expected for PB-mediated integration (Yoshida et al. 2017) (Fig. 1B-C and S1C). Within gene bodies, integrations were more frequent near gene 5’ ends, with this bias further accentuated for barcodes with detectable expression (Fig. S1D). After normalising for the number of available TTAA sites in each gene, essential genes showed a higher PB integration density than non-essential genes. This enrichment was largely attributed to essential genes being more likely to be expressed, although we still observe a modest effect when restricting the analysis to expressed genes (Fig. S1E). Of the 19,120 human protein-coding genes in the MANE annotation (Morales et al. 2022), 5,095 (Regular) and 5,015 (PCPA) had at least one integration somewhere within the gene body. Of these, 508 and 478, respectively, had at least one integration in an exon.

We also generated two matched reporters in which transcription was driven by the constitutive, moderately active PGK promoter (Qin et al. 2010) (Fig. S1F). In these reporters, the promoter was followed by the first exon and intron of β-globin to promote engagement of the splicing machinery and more closely resemble a conventional transcription unit, and then the sequences matching the “Regular” and “PCPA” reporters. We mapped 1,955 and 3,320 integrations of these reporters, which were distributed similarly to those of the promoterless reporters (Fig. S1G and Table S3). As expected, promoter-driven reporters were expressed at most integration sites and exhibited comparatively limited variation in expression (Fig. 1D). By contrast, promoterless reporters were expressed at only ∼40% of integration sites, consistent with more dependency on integration into endogenous, actively transcribed regions.

### Reporter exons are typically spliced to host exons

We next asked whether the reporter splice donor engages downstream host splice acceptors in endogenous host transcripts. To capture host transcript sequences immediately downstream of the β-globin sequence in the reporter, we adapted the genomic mapping strategy, using DpnII restriction of cDNA followed by inverse PCR, an approach we will refer to as ‘reporter-host connectivity mapping’ (Fig. 1E). If splicing between the reporter exon and the downstream host exon occurs, we expect the paired-end reads of these mapping libraries to capture the sequence between the next exon and the subsequent DpnII restriction site in the processed RNA. Conversely, if splicing does not happen, we expect to obtain the sequence downstream of the β-globin exon in the reporter up to the nearest DpnII recognition site in the genome. We obtained at least 5 usable reads for 6,385 and 6,068 unique barcodes in the Regular and PCPA libraries, respectively, >40% of which were barcodes mapped to MANE introns. For each reporter, we classified the cDNA products based on the read-mapping pattern, finding that most barcodes were associated with a specific 3’ splicing pattern (see Methods, Fig. S2A).

In most cases, integrations into MANE introns produced the expected splicing pattern from the reporter to the subsequent host exon (Fig. 1F-G). We further validated two of these splicing events by RT-PCR and Sanger sequencing (Fig. S2B). The rarer exonic integrations more often resulted in no downstream splicing, whereas integrations outside of annotated gene bodies were also often spliced. When splicing was not to the subsequent annotated exon, it was to other exons annotated in MANE or GENCODE, or to sequences with reported autonomous exon activity in a plasmid-based reporter assay that profiled the whole human genome (Stepankiw et al. 2023) (Fig. 1H). Overall, considering both reporters, the introduced sequences underwent splicing at 3,702 distinct loci, supporting their ability to undergo splicing as an autonomous units. We also observed rare splicing events to cryptic exonic sequences within the reporter, including in the PB 3’ ITR and in the GNAI1 intronic sequence in the PCPA vector (Fig. S2A and S2C).

Next, we adapted our method to capture splicing upstream into the 5’ acceptor site in the reporter (Fig. S2D). We were able to validate splicing from the upstream exon to the reporter for several integrations (Fig. S2E). This analysis revealed that the PB ITRs contain a cryptic exon that was commonly used, producing a consistent 75 bp upstream exon included before the β-globin exon of the reporter (Fig. S2C). Although we recovered several transcripts spliced to upstream host exons, large-scale analysis of these events as above was not feasible. The β-globin exon upstream of the barcode is only 53 bp, leaving insufficient sequence for the required two rounds of nested inverse PCR. These chimeric transcripts could therefore only be identified in individual reads. Below, we performed more in-depth analysis of the upstream splicing pattern for the two-exon reporters.

To further elucidate RNA processing at each integration site, we performed oligo-dT–based poly(A) selection and quantified barcode abundance in both the Poly(A)+ and Poly(A)-fractions; additionally, we made libraries from amplicons that spanned the ß-Globin exon and the downstream GNAI1 intron and, for the PCPA reporter, the region downstream of the PCPA site. Across the majority of integrations, barcodes were depleted from the Poly(A)− fraction and the downstream intronic amplicon relative to Poly(A)+ and total expression, consistent with efficient splicing and downstream polyadenylation of the reporter transcript irrespective of integration context (Fig. S2F). Depletion from the Poly(A)− fraction was most pronounced among the highest-expressed integrations, which were enriched for expressed genomic regions, active transcription-associated chromatin states, and proximity to CAGE clusters.

Insertion of TRIP reporters into endogenous transcripts will often disrupt open reading frames (ORFs), generating frameshifts and downstream premature termination codons (PTCs), potentially making the chimeric transcripts susceptible to nonsense-mediated decay (NMD). To assess this, we inhibited NMD using cycloheximide (CHX) to arrest translation, essential for NMD activity, and quantified changes in reporter expression relative to vehicle-treated controls (EtOH). Reporters predicted to trigger NMD by inducing a PTC more than 55 bp upstream of the last exon-junction complex in the full transcript were strongly stabilized following CHX treatment in both the Regular and PCPA libraries, as expected. In contrast, reporters predicted not to trigger NMD, or those integrated outside MANE genes, as a group not affected by translation inhibition (Fig. 2A). Consistent with the simple 50-55 nucleotide NMD rule (Kurosaki et al. 2019), we observe that transcripts expected to contain a PTC once the reporter splices into host transcripts were equally affected by NMD inhibition (Fig. S3A). The degree of stabilization depended on position, with integrations near the beginning of transcription units showing no stabilization (Fig. S3B), consistent with observations that PTCs close to the initiation codon AUG evade NMD (Romão et al. 2000; Peixeiro et al. 2012). Together, these results establish that promoterless TRIP reporters are incorporated into endogenous transcripts and engage the surrounding RNA-processing machinery.

**Figure 2.**
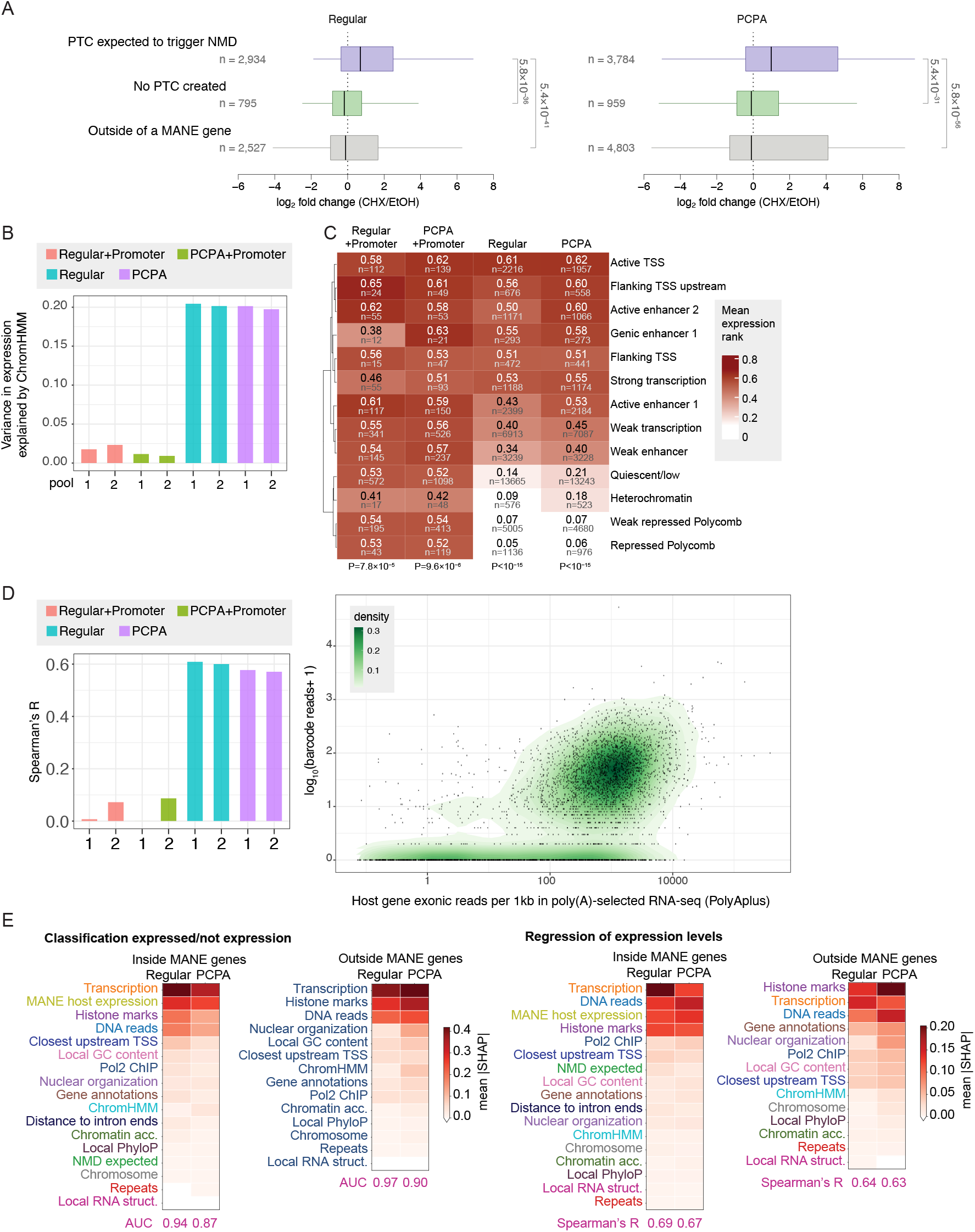
TRIP reporter exons expression depends on host gene context. **(A)** Changes in expression of mapped barcodes upon arrest of translation using cycloheximide (CHX) for 4 hours. Barcodes are divided based on their presence within a MANE transcript and the potential to trigger NMD according to the currently accepted model (see Methods). **(B)** Fraction of the variance in log_10_-transformed read counts explained by the ChromHMM state of the barcode position. **(C)** Mean barcode expression ranks (1 - most abundant barcode; 0 - least abundant, barcodes with identical values are assigned the minimal rank) for barcodes mapped to different ChromHMM states. **(D)** Left: Correlation between host gene expression and barcode expression for different reporter types; Right: Correlation between barcode expression in the regular TRIP pool and host gene expression. Host gene expression taken from the ENCODE K562 polyA-selected RNA-seq data. **(E)** SHAP values for the indicated sets of features for held-out data, for XGBoost classifier trained to predict if a barcode has any expression reads (left) or for predicting the actual log10-transformed read count (right). The ROC AUC and Spearman R of the classifiers are indicated below each heatmap.

### Promoterless reporter expression is shaped by the surrounding genomic environment

To identify genomic features associated with reporter expression, we linked barcode RNA abundance to the corresponding integration site and examined the relationship between local chromatin properties and expression. Consistent with their overall higher activity, reporters carrying an intrinsic promoter were robustly expressed across diverse genomic environments with little variation. Chromatin state only explains ∼2% of the variation in expression (Fig. 2B), and mean expression rank varied over a narrow range across all thirteen ChromHMM states (Fig. 2C). Integrations into heterochromatic regions were generally expressed at the lowest level, consistent with earlier TRIP studies using different intrinsic promoter-containing constructs (Brueckner et al. 2016; Akhtar et al. 2013).

Promoterless reporters, by contrast, were strongly sensitive to their local chromatin environment (Fig 2B). They were expressed at low levels when integrated into repressive chromatin states such as heterochromatin, Polycomb-repressed, and quiescent states (Fig. 2C). In contrast, they were robustly expressed within transcribed regions, thus recapitulating the expression behavior of the native surrounding endogenous genes. Notably, expression was not a uniform feature of active chromatin; among transcriptionally active states, promoterless reporter expression was highest at transcription start sites and enhancers and comparatively lower within gene bodies. Enhancers are sites of high RNA Pol II activity that generate enhancer RNAs, which are quickly turned over (Field and Adelman 2020). We therefore hypothezise that integration in these regions could capture otherwise unproductive transcriptional events, and the reporter-embedded splice sites can trigger splicing into the second exon of a downstream gene, thereby generating stable transcripts. Consistent with this model, our reporter-host connectivity mapping data show integrations in enhancers giving rise to chimeric transcripts with the downstream gene. The length of the 3′ intron and the addition of the PCPA sequence did not measurably alter overall expression sensitivity to chromatin state. Reporter expression further correlated with the endogenous gene expression at each integration site, confirming that the promoterless reporter quantitatively reflects the productive transcriptional activity of its local environment (Fig. 2D). In contrast, reporters carrying an intrinsic promoter sequence do not show this correlation, as expected (Fig. 2D). To obtain a more global view of how different features of the integration site shape reporter expression, we trained an XGBoost model using both global features, such as overlap with exons or introns of MANE or GENCODE annotations; host gene expression; low-resolution features such as TSA-seq derived distance to nuclear speckles; as well as local features, such as nascent RNA-seq coverage, histone marks, and GC-content in a 100 nt window centred at the integration site; and the number of reads obtained from the DNA libraries, reporting on the relative copy number of the reporter in the cells. The models could predict with reasonable accuracy both whether a reporter was expressed, i.e., had at least one expression read, both for barcodes integrated inside MANE gene bodies and for those outside, with AUC of 0.94–0.97 and 0.87–0.9 for the Regular and PCPA reporters, respectively and the expression levels (Spearman’s R of 0.63–0.67, Fig. 2E). SHAP analysis of the feature groups that contributed the most to model predictions showed that transcription and histone marks had the highest contribution to model accuracy, with the expected contribution of DNA copy number, with the proximity to upstream TSS having an independent contribution, consistent with the higher expression observed in promoter-proximal states (Fig. 2E).

### A PCPA site is rarely used outside of its native context

Having established the integration landscape, we next investigated whether the cryptic polyadenylation site located within the 3’ region of our PCPA reporter undergoes CPA upon U1 treatment. In its endogenous context, U1 snRNP actively suppresses this site in the *GNAI1* intron in HeLa cells (Fig. S1A). To test whether this sensitivity varies by genomic position, we treated the TRIP pools with U1 antisense morpholino oligonucleotides (AMOs) that disrupt the U1 snRNP (Kaida et al. 2010). We confirmed successful disruption of U1 snRNP using RNA-seq, which revealed the expected pattern of increased intron retention and premature termination in the first introns of known U1-sensitive genes (Oh et al. 2017), as well as missplicing of a previously reported U1-sensitive transcript (Berg et al. 2012) (Fig. S3C-D).

To quantify the effects of U1 inhibition on the PCPA reporter, we performed PolyA selection followed by reverse transcription using primers positioned either within the ß-globin exon (‘exonic’) or upstream or downstream of the PCPA site and quantified barcode expression (Fig. 3A). In the sites where the PCPA site becomes active upon U1 treatment, we would expect to see a preferential reduction in amplicon reads from the amplicon using the downstream primer. The changes in barcode expression upon U1 treatment were highly correlated between the two RT primers (Spearman R>0.9), suggesting that there is no strong PCPA induction. We then stratified the barcodes based on the ChromHMM state, and found that barcodes integrated into promoter-proximal regions were upregulated upon U1 treatment (Fig. 3B), consistent with the reported changes in RNA Pol2 occupancy for promoter-proximal regions (Mimoso and Adelman 2023) and increased expression of short genes upon U1 treatment (Oh et al. 2017).

**Figure 3.**
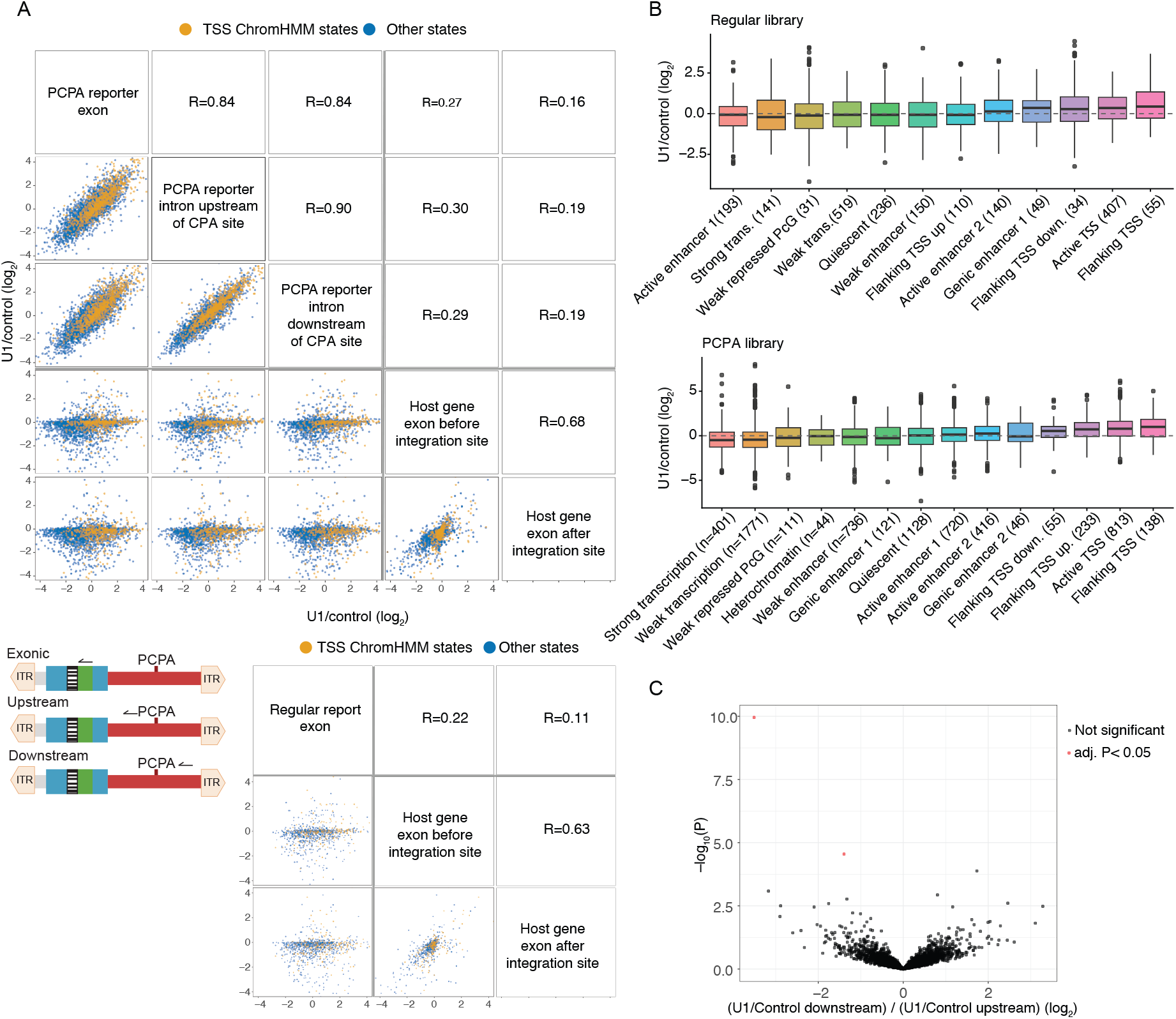
A randomly integrated PCPA-sensitive site rarely undergoes PCPA. **(A)** Correlations between changes in expression of barcodes as measured by amplicon sequencing using three different RT primers (illustrated at the bottom left graphic), and expression changes in host exons immediately preceding or succeeding the barcode location. In each scatter plot, barcodes mapped to ChromHMM TSS-proximal states are in orange. Spearman correlation coefficients are shown. The bottom plot compares barcode and host exon expression for the Regular library, where only regular exonic expression was measured. **(B)** Changes in barcode expression (exon amplicons) as a function of the ChromHMM state it maps to. **(C)** Differential expression of barcodes upon U1 treatment as computed by DESeq2, comparing changes in expression using downstream RT primer to an RT primer upstream of the GNAI1 PCPA site.

Within gene bodies, different integration contexts can change barcode expression. First, if the barcode is integrated downstream of a PCPA site in the host gene, we expect reduced barcode expression regardless of the RT primer used, as transcription of the host gene is generally expected to no longer reach the barcode. Indeed, changes in barcode expression, in particular for the barcodes repressed by U1, were correlated with expression changes of the host exon preceding the barcode (Fig. 3A). When we used DESeq2 to specifically compare changes in expression as measured using primers downstream vs. upstream of the CPA site, and found that only two barcodes showed reduced expression specifically downstream (adjusted P<0.05, Fig. 3C), which would occur if the CPA site was used upon U1 treatment. We conclude that a sequence that can effectively drive PCPA at its endogenous locus fails to do so at thousands of other genomic contexts.

### Partly retained introns preserve splicing behavior outside their native genomic context

The strong context dependence of PCPA site usage prompted us to ask whether other co-transcriptional RNA-processing events are similarly dictated by genomic position. We therefore considered partially retained introns, whose splicing could in principle be determined either by intrinsic features of the intron or by properties of their endogenous transcriptional and nuclear environment. Previous systematic analyses have shown that distinct classes of retained introns are associated with specific nuclear domains (Barutcu et al. 2022), including nuclear speckles and the lamina. We therefore developed IntronTRIP – a TRIP-based intron reporter to systematically determine how genomic integration site and host-gene context influence intron splicing. We initially cloned five speckle-associated introns, five lamina-associated introns, and five constitutively spliced introns, together with 50 bp of their endogenous flanking exonic sequences, nested into another 50 bp from the ends of β-globin exon 2. These reporters have the same 5’ 64 bp ß-Globin intron 1 sequence as for single exon TRIP. To avoid mixing two different classes of special intronic sequences, the *GNAI1* intron with its PCPA site at the 3’ end of the reporter was replaced with the first 50 bp of the second ß-Globin intron (Fig. 4A). Following random integration into K562, we assessed the global splicing behavior of all 15 introns. All five constitutive introns were overall efficiently spliced, all nuclear speckle-associated introns were spliced at intermediate levels, while of the five lamina-associated introns one was not efficiently spliced and the other four were spliced quite efficiently (Fig. S4A). Lamina-associated retained introns are generally longer than speckle-associated ones, so comparing the two classes directly would confound intron class with intron length. We therefore selected lamina-associated introns of comparable length to the speckle-associated set. These generally exhibit lower endogenous retention than the longer introns typical of lamina-associated introns. Because most lamina-associated introns and all constitutively spliced introns behaved similarly and showed little evidence of splicing variation, we selected to continue with two introns partially retained and associated with the lamina in their endogenous loci, from *PACRGL* and *TTC17*, as well as the five speckle-associated introns, from *NBEAL2*, *TONSL*, *USP21*, *KPTN*, and *DDX51*. As a control, we included the constitutively spliced intron from *PACRGL* that shows no evidence of intron retention. Overall, we generated TRIP reporters from eight introns of lengths 126–509 bp (Fig. 4B).

**Figure 4.**
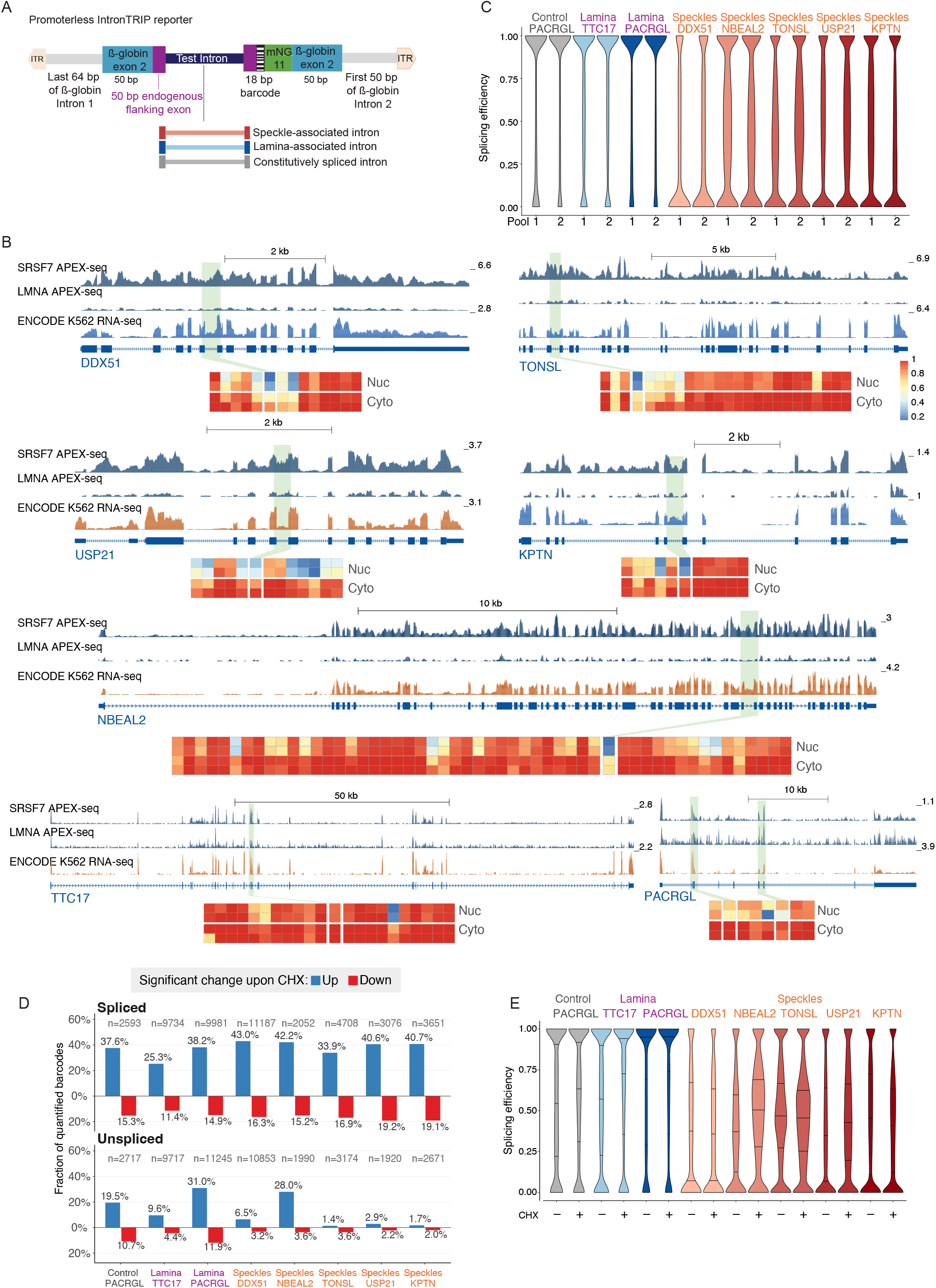
IntronTRIP integration mapping. **(A)** Outline of the IntronTRIP approach. **(B)** The eight introns used in IntronTRIP assays. For each intron, the APEX-seq coverage using a speckle marker (SRSF7) and a lamina marker (LMNA) from (Barutcu et al. 2022) are shown, normalized to the same scale. RNA-seq from K562 cells from ENCODE (PolyA-selected libraries) is also shown, with red representing (+) strand and blue (-) strand. The introns used in IntronTRIP are highlighted. For PACRGL, the first shaded intron is the lamina-associated one, and the second shaded one is the intron used as a control. Below each intron, the heatmap shows average splicing efficiencies in the Nuclear and Cytoplasmic fractions (two replicates each), obtained from ENCODE fractionation RNA-seq data in K562 cells. **(C)** Distributions of splicing efficiencies – “spliced reads / (spliced & unspliced reads)” for the indicated introns in each of the two IntronTRIP pools obtained for each intron. **(D)** Fraction of barcodes in the indicated IntronTRIP pools that was significantly upregulated (blue) or downregulated (red) upon NMD inhibition, in the spliced isoform (top) or the unspliced isoform (bottom). Significance was determined using DESeq2. n indicates the number of barcodes for which the respective isoform could be quantified. **(E)** Distribution of splicing efficiencies for the indicated introns after 4h of CHX treatment (+) or 4h treatment with the vehicle control ethanol (-).

Following random integration into K562 cells, we mapped reporter integration sites and mapped 50,831–268,899 integrations per intron, with two pools per intron (Table S4), which spanned all chromatin states (Fig. S4B). Integration and expression patterns were consistent across the eight IntronTRIP pools, with 38–59% of integrations being expressed (Fig. S4C). Expression behavior depended on very similar features as those observed from single exon integrations, and could be predicted from genomic features with similar accuracy; the strongest contribution being host gene expression, with additional contributions from chromatin state, DNA copy number, and histone marks (Fig. S4D-E).

We validated that the IntronTRIP reporters were integrated into host transcripts. Similar to single-exon TRIP, intronic integrations produce the expected splicing to the downstream exon in >75% of the cases, and such mapping was validated for 1,507 barcodes across the five introns for which we performed such analyses (Fig. S5A). For three of the intron reporters, mapping upstream splicing further confirmed the expected spliced pattern; integrations into MANE introns were mainly spliced to the previous MANE exon (Fig. S5B). For some integrations, we were able to confirm splicing to both upstream and downstream exons (Fig. S5C). The few integrations that mapped to exons, mostly last exons, were much less efficiently spliced, consistent with efficient co-transcriptional splicing of most conventional introns (Reimer et al. 2021).

We next quantified reporter splicing at each integration site using three barcode-specific amplicons that selectively captured spliced, unspliced, or total reporter transcripts. To focus on mature RNA products, RNA was enriched for polyadenylated transcripts prior to reverse transcription. Splicing, requiring at least 20 reads from both spliced and unspliced amplicons, could be quantified for 19,606–73,981 unique barcodes per intron, across 19,016-67,475 unique positions (Table S4). Strikingly, despite integration across diverse genomic environments, the five partially-retained speckle-associated introns largely preserved their characteristic endogenous splicing behavior. In contrast, the two lamina-associated introns and the constitutively spliced control were efficiently spliced across nearly all integration sites (Fig. 4C).

Due to their larger total exonic length, IntronTRIP reporters were expected to lead to NMD more often than single integrations. Indeed, upon treatment with CHX, we observed preferential up-regulation of integrations inside MANE introns, with spliced introns up-regulated consistently across all eight introns and with similar effect sizes (Fig. 4D and S5D-E; Table S5) of 1.6-2.5 log2-transformed fold-change in the spliced isoform (Fig. S5E), and overall integrations predicted to cause NMD were preferentially induced (Fig. S5D). Similarly to single exon integrations, NMD was preferentially suppressed for integrations close to the 5’ ends of transcripts (Fig. S5E). When comparing control and CHX-treated cells, the similar effect sizes observed across all eight introns led to an overall similar distribution of splicing efficiencies between control and CHX-treated cells (Fig. 4E). Therefore, below we interpret the distribution of splicing efficiencies between introns at different genomic positions as consequences of differential splicing efficiencies rather than differential sensitivities to NMD.

Although all five speckle-associated introns retained partial splicing across genomic locations, they differed in how splicing was distributed between positions. The DDX51 was the most extreme of the five, with individual integrations tending toward either efficient or minimal splicing and the fewest intermediate values (Fig. 4C). As shown below, DDX51 is also the intron whose splicing depends most strongly on host gene expression and on proximity to nuclear speckles, indicating that its processing is the most sensitive of the five to particular features of its genomic environment.

### Genomic and sequence correlates of splicing efficiency

Having established that partly retained introns maintain their overall splicing behavior across thousands of genomic positions, we next asked how sensitive their splicing is to the local genomic context. When comparing ChromHMM states (Fig. 5A), we found that the five speckle-associated introns all behaved differently from the two lamina-associated introns and the constitutively spliced one, in that their splicing was less efficient in regions associated with open chromatin and in regions with promoter/enhancer activity, but also in regions associated with silent chromatin (which were generally depleted of expressed barcodes). We note that the reduced splicing efficiency in these states could also reflect less efficient exon recognition and splicing into host transcripts, rather than differential sensitivity of the introns themselves (see Discussion). By contrast, in transcribed regions away from promoters and enhancers (‘strong transcription’ state), all eight introns reached similar mean splicing efficiencies, indicating that the differences between intron classes are enhanced in particular chromatin states rather than being uniform across the genome. While most of the speckle-associated introns behaved similarly, DDX51 was evidently more sensitive to highly active chromatin regions (Fig. 5A).

**Figure 5.**
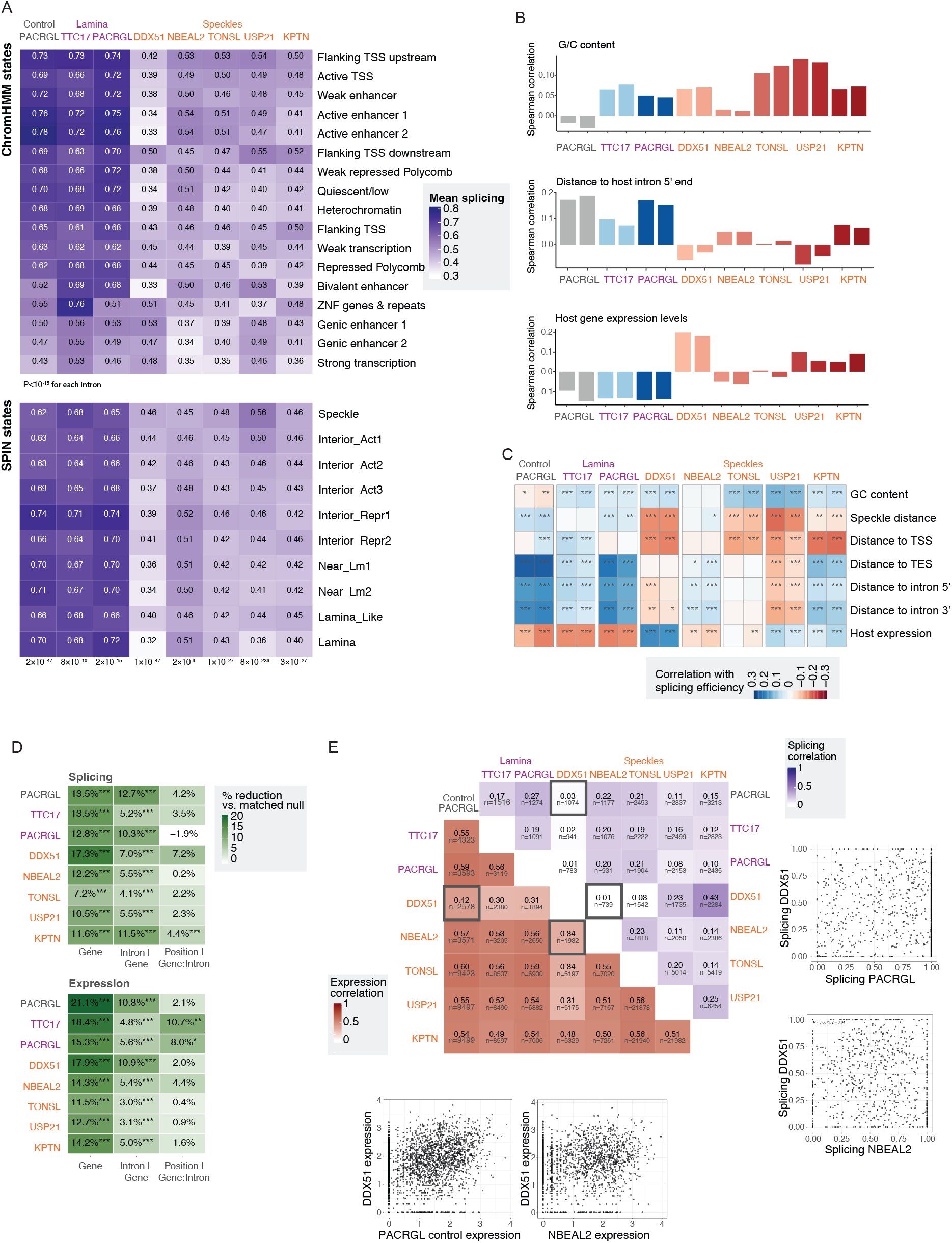
Determinants of different introns’ splicing efficiency. **(A)** Mean splicing efficiency for reporters of the indicated introns, stratified by ChromHMM state (top) and SPIN state (bottom) of the barcode position. Means of the two pools for each intron are shown. Kruskal-Wallis P-values for the influence of the state on splicing are shown in each heatmap. **(B)** Spearman correlation coefficients between splicing efficiency and the indicated feature. **(C)** Spearman correlation coefficients and significance for the indicated features. TSS, TES, intron, and host expression correlations are computed relative to MANE host genes, only for barcodes hosted in MANE genes. Speckle distance is from TSA-seq (Y. Chen et al. 2018). **(D)** Difference between barcodes at the indicated level compared to barcodes at the previous level. % reduction is the percentage by which the difference is smaller for barcodes that share the level than for a randomly chosen, equally-sized set of barcodes drawn from the appropriate comparison pool (random barcode pairs for the same gene; the same gene, for the intron-level test; the same intron, for the position-level test). A % reduction near 0% means sharing that level carries no extra information — those barcodes look just as different from each other as random ones would. Higher percentages mean barcodes sharing that level are progressively more similar to one another than chance predicts. Significance was assessed empirically by directly comparing to 8000 random resamplings. **(E)** Correlation coefficients between expression levels (lower triangle) and splicing efficiencies (upper triangle) for pairs of IntronTRIP reporters integrated by chance into the same positions. Two scatter plots are shown, and the corresponding squares are outlined with thick borders in the heatmaps.

We next examined the effects of spatial location of the locus in the nucleus. We used the 10 SPIN states, defined based on integrating data from TSA-seq, which measures proximity of genomic regions to nuclear speckles, and Hi-C data that measures proximities between genomic regions (Y. Wang et al. 2021) (Fig. 5A). Here, there was less variability between states, but interestingly, different speckle-associated introns had opposing trends, with DDX51, and to a lesser extent TONSL, USP21, and KPTN splicing better when placed in speckle-proximal regions, and NBEAL2 in contrast being more efficient near the lamina. Notably, the greater speckle-proximity sensitivity of the DDX51 intron relative to the NBEAL2 intron mirrors the relationship at their endogenous host loci: while both are speckle-associated, DDX51 lies closer to nuclear speckles than NBEAL2, with SON TSA-score in the 97th and 95th percentiles, respectively (Y. Chen et al. 2018).

We then examined other features (Fig. 5B-C) and found variable correlations with splicing efficiency, proximity to the preceding host 5’ end (considering only barcodes inside MANE introns), and host gene expression, with most correlations being statistically significant and often in opposing directions for different introns (Fig. 5C). When combined into XGBoost predictive models, splicing efficiency could be predicted from genomic annotations and local features with Spearman R of 0.29 and 0.31 for integrations inside and outside of MANE genes, respectively (Fig. S4D). This accuracy was notably lower than what could be achieved with expression, but splicing efficiency estimates were also substantially noisier than estimates of expression, likely due to an inherently lower dynamic range. When considering different sets of genomic features, overlap and distances to gene annotations, including exons, transcription levels, and histone marks, contributed the most to the prediction accuracy, with notable contribution of nuclear organization, particularly distance of the locus to nuclear speckles (Fig. S5F).

To obtain a more holistic view of the influence of local environment on expression and splicing efficiency, we compared integrations sharing progressively more local genomic environments. We compared the reduction in variability between barcode pairs if they were co-hosted in the same gene, same intron or same position (Fig. 5D), evaluating significance empirically by comparing to thousands of random resamplings. Host genes typically had a substantially higher effect on expression levels than on splicing, with the notable exception of DDX51, consistently with its splicing being most influenced by host expression levels. For the other seven introns, the effects of the specific host intron (after controlling for the host) were larger for splicing than for expression, and for all introns this effect was significant. For a subset of introns, there was also a significant effect for the specific integration position within the host intron. Thus, splicing and expression appear to respond to genomic context at different spatial scales; expression is more associated with the broader host-gene environment, whereas splicing is more sensitive to more local features surrounding the integration site, yet different introns are affected by different features, and to varying degrees.

### Direct comparison of integrations at shared positions reproduces genome-wide patterns

A limitation of the TRIP approach is that each intron was integrated in a different set of positions, so while robust and differential trends were observed, we also wanted to compare head-to-head different introns integrated in shared positions. We first evaluated different IntronTRIP reporters integrated at the exact same genomic positions. Such overlaps were observed more often than expected by chance; for example, the NBEAL2 and DDX51 reporters were each integrated into 0.26–0.34% of the genomic TTAAs, yet preferential integration into open chromatin resulted in overlaps at 1,938 positions where expression could be measured (11.6-fold more than expected by chance) and 739 shared positions with sufficient reads for measuring splicing (36.3-fold enrichment) (Fig. S6A). Across all 28 pairs of introns, expression measured at the same position correlated strongly for the 21 pairs not involving DDX51 (Spearman R 0.58 to 0.60), yet splicing correlated weakly (Spearman R= −0.03 to 0.43). DDX51 was a notable outlier, as its expression correlated little with other introns, associated with its overall broader expression, in particular from regions with weak enhancer chromatin states (Fig. 5E and Fig S6C-E). It is possible that this intron has the ability to boost transcription from weakly transcribed regions, as described for some introns (Uriostegui-Arcos et al. 2023) (see Discussion). Introns of the same class did not exhibit higher splicing correlation than introns of different classes, the single exception being DDX51 and KPTN. Of the five speckle-associated introns, DDX51 and KPTN combine high intronic GC-content with relatively lower exon-intron GC contrast, known to influence intron retention (Tammer et al. 2022); this may suggest the relatively better correlation of the DDX51 and KPTN pair at shared positions and stronger speckle proximity dependency.

Shared positions were strongly biased toward open chromatin near transcription start sites and enhancers (Fig. S6A). To mitigate this, we extended our approach and used a two-step TRIP to first integrate a landing pad at many different locations, and then recombine our intron reporter into the established landing pad, similar to what has recently been used for enhancers and insulators (Hong et al. 2024; Y. Wu et al. 2025). We integrated a landing pad carrying the reporter gene TagBFP flanked by the attP sites of the two serine recombinases φC31 Bxb1 for efficient directional recombination (Fig. S6B) (Xu et al. 2013). Each landing pad carries a unique barcode for identification of the genomic location (gBC). By FACS of TagBFP-positive cells, we generated four pools carrying landing pads. The introns from DDX51 and NBEAL2 showed the most different sensitivity to both nuclear localization and local genomic context (Fig. 5A-B). We, therefore, chose these two and recombined them into the TRIP landing pad pools.

We mapped 130,832 landing pad sites with unique barcodes (Table S6), and NBEAL2 and DDX51 reporters were recombined and detected at 20,683 and 27,980 positions, and recapitulated the main trends from the original IntronTRIP libraries (Fig. S6C). While the landing pad approach did not increase the number of NBEAL2/DDX51 co-integration positions (2,303 for regular IntronTRIP vs. 2,138 for landing pad), these positions were substantially less biased toward promoter- and enhancer-associated states (Fig. S6A, see Discussion). The splicing efficiencies of the two introns were generally correlated less than expression levels (Fig. S6E), and for the landing pad integrations, the changes in splicing were decoupled from changes in expression (Fig. S6F). Consistent effects were observed when stratifying the shared positions by ChromHMM states – NBEAL2 was spliced better than DDX51 in enhancer-associated states – and DDX51 splicing was generally more efficient than that of NBEAL2 in GC-rich regions (Fig. S6G). When considering SPIN states, DDX51 was consistently spliced better than NBEAL2 preferentially in speckle-proximal and interior active states (Fig. S6G). The splicing of partly-retained speckle-associated introns thus responds differentially to the genomic and nuclear environment in a manner dependent on their sequences.

### Genomic context modulates the SON dependence of speckle-associated intron splicing

To further investigate how genomic context influences the splicing of nuclear speckle-associated partly retained introns, we perturbed nuclear speckle function through SON depletion. SON and SRRM2 form the structural core of nuclear speckles, with SON appearing to have a particularly important role in maintaining speckle function, as depletion of SON alone largely recapitulates the effects of combined SON and SRRM2 depletion (Małszycki et al. 2026). Because prolonged SON depletion can cause mitotic arrest and other secondary effects (Ahn et al. 2011), we generated a K562 cell line in which endogenous SON was Halo-tagged, enabling rapid depletion of Halo-SON and allowing us to examine its immediate effects on splicing (Fig. S7A). We validated functional SON depletion by RT-PCR analysis of known SON-sensitive splicing events (Fig. S7B) and then performed RNA-seq upon SON depletion to assess transcriptome-wide changes in RNA processing. Because our TRIP libraries included introns selected from nuclear speckle-associated partly retained transcripts, we next examined whether the corresponding endogenous introns responded to SON depletion in their native genomic context. Consistent with their proximity to nuclear speckles, depletion of SON leads to increased intron retention for those of *NBEAL2*, *TONSL*, and *DDX51*, while the *USP21* and *KPTN* introns are less affected (Fig. 6A, S7C), similar to what was shown in an HCT116 cell line (Małszycki et al. 2026). As only a subset of introns require the activity of SON and nuclear speckle’s integrity for their splicing, the constitutively spliced intron from *PACRGL* was not affected after SON depletion (Fig. 6A). In our HaloTag-SON K562 line, we generated TRIP pools of the *NBEAL2*, *TONSL*, *DDX51*, and control *PACRGL* introns to investigate how genomic context affects dependency on functional nuclear speckles for splicing. We validated successful integrations as described above with 30,820–69,723 barcodes into 16,381–26,350 unique genomic positions (Table S7), and sequenced RNA amplicons from lines treated with DMSO vehicle or HaloPROTAC3.

**Figure 6.**
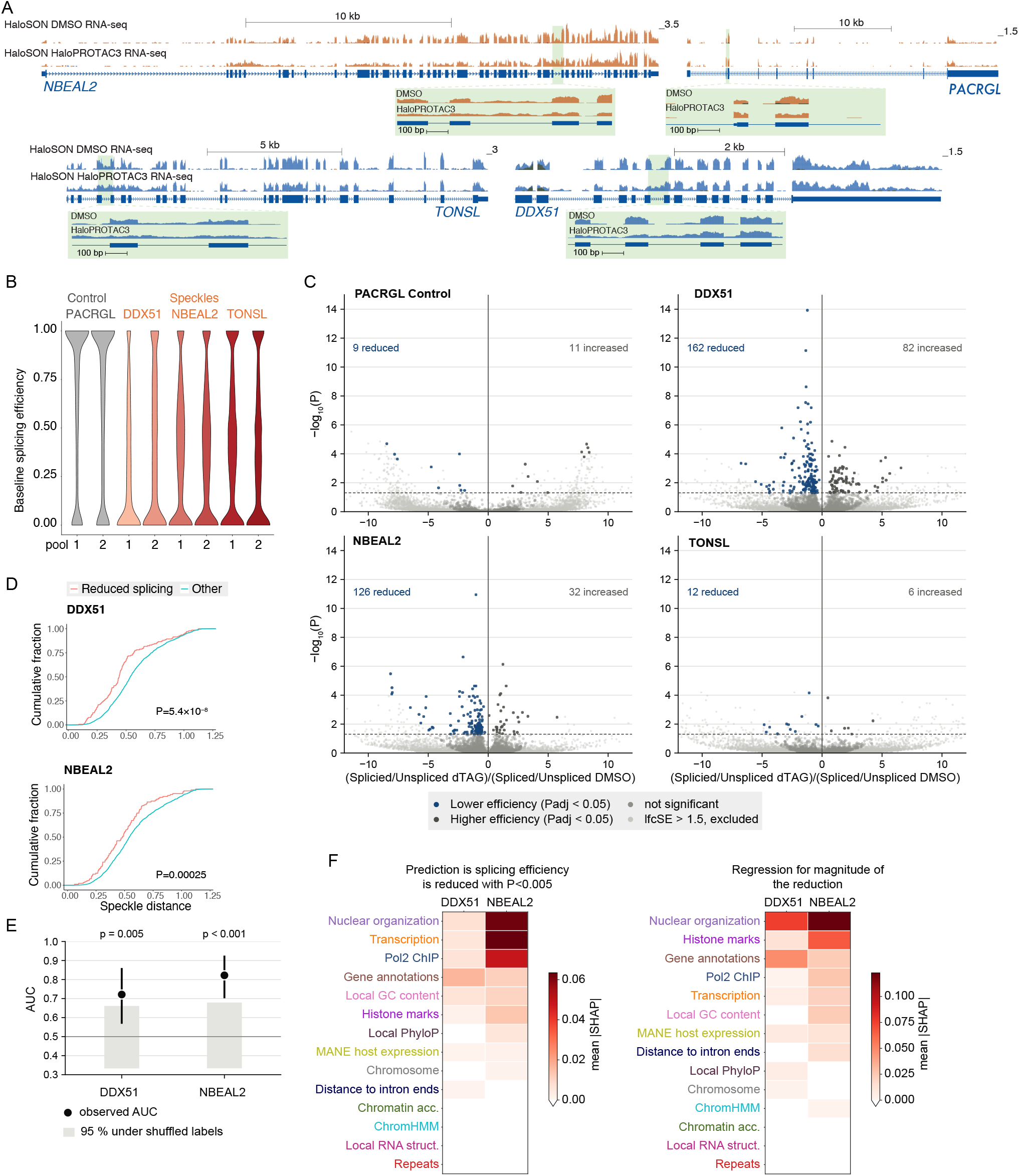
Changes in reporter splicing efficiency upon depletion of SON. **(A)** RNA-seq coverage on the original genes from which reporter introns were taken upon SON depletion in K562 cells. 1 kb zoom-in on the TRIP introns are shown below. **(B)** Splicing efficiencies of the indicated introns in control DMSO-treated K562 HaloTag-SON cells. **(C)** Changes in splicing efficiencies for the indicated IntronTRIP reporters as evaluated by DESeq2 comparing the numbers of spliced and unspliced reads between DMSO- and HaloPROTAC3-treated cells. **(D)** Distances to speckles of the barcodes integrated in MANE genes from C. For each barcode, the average distance of the host MANE gene is used. P-values are computed using the Wilcoxon rank-sum test. **(E)** Performance of XGBoost models trained on real vs. shuffled data. Empirical p-values are shown. **(F)** SHAP values for the indicated feature groups for classifiers trained to differentiate between barcodes with reduced splicing upon SON depletion vs. other barcodes with lfcSE<1.5 (left), and a separate classifier trained to predict the magnitude of the reduction in splicing efficiency (right).

In DMSO-treated cells, intron splicing efficiencies closely recapitulated their dependencies observed with original integrations into unedited cells, highlighting the robustness of our method (Fig. 6B and S7D-E). We then treated cells with either HaloPROTAC3 or the vehicle control DMSO in triplicate and used DESeq2 to quantify the effects on splicing. For the control *PACRGL* intron and the speckle-associated TONSL intron, relatively few integrated introns were significantly affected, whereas hundreds of integrations of DDX51 and NBEAL2 exhibited reduced splicing (Fig. 6C). The affected integrations were located in genes that are natively located closer to speckles than other integrations (Fig. 6D).

To evaluate the features predictive of sensitivity to SON levels, we built XGBoost classifiers to predict whether insertions of DDX51 and NBEAL2 undergo less efficient splicing upon SON depletion with unadjusted P<0.005 upon SON depletion (one classifier per intron) and separate classifiers for predicting the magnitude of the effects on the affected introns. The prediction classifiers obtained a significant AUC of 0.72 and 0.82, and those trained to predict the level of reduction obtained Spearman R of 0.38 and 0.37 (Fig. 6E). Examining the features supporting the predictive power of these classifiers revealed that while proximity to speckles had a substantial contribution for both introns, the features predictive of changes in splicing of DDX51 and NBEAL2 were often different (Fig. 6F), consistent with the notion that while SON presence and speckle integrity are instrumental for the proper splicing of both introns, the effects of splicing loss are further modulated by the local genomic context.

### Intron identity sets the export fate of the host transcript

Intron retention is widespread in mammals and regulates gene expression through NMD and nuclear sequestration (Braunschweig et al. 2014). Many speckle-associated retained introns are also within the previously identified class of detained introns, and are therefore thought to act, in part, by restricting export from the nucleus (Barutcu et al. 2022; Boutz et al. 2015); although some transcripts with retained introns do reach the cytoplasm (Monteuuis et al. 2019; Y. Li et al. 2006). In our intron set, the two introns from *NBEAL2* and *DDX51* have been characterized as detained introns (Boutz et al. 2015). We therefore investigated the splicing state of the introns in different cellular fractions by fractionating IntronTRIP cell pools. We confirmed successful fractionation of K562 cells by strong nuclear enrichment of MALAT1 and cytoplasmic enrichment of ACTB (Fig. S8).

In the nuclear fraction, we found for all introns that most barcodes produced a mixed state with both spliced and unspliced introns (Fig. 7). In contrast, in the cytosol there was a substantial difference between the introns (Fig. 7). Transcripts containing the lamina-associated or control intron were essentially fully spliced in the cytoplasmic fraction, whereas most speckle-associated introns, with the exception of the one from KPTN, contained a major portion of intron-containing molecules in the cytosol (Fig. 7). The same pattern was observed in the endogenous genes: in K562 fractionation RNA-seq, these introns were retained in the cytoplasm relative to other introns of the same genes, whereas the lamina-associated, control, and KPTN introns were not (Fig. 4B). Overall, this shows that intron-retaining transcripts from most nuclear speckle introns reach the cytoplasm at ectopic positions, indicating that these introns do not act as absolute nuclear-retention signals. Intron identity nonetheless determines the fate of the transcript it sits in, transcripts carrying partially-spliced introns are exported across thousands of ectopic positions, similar to the case at the introns’ native loci.

**Figure 7.**
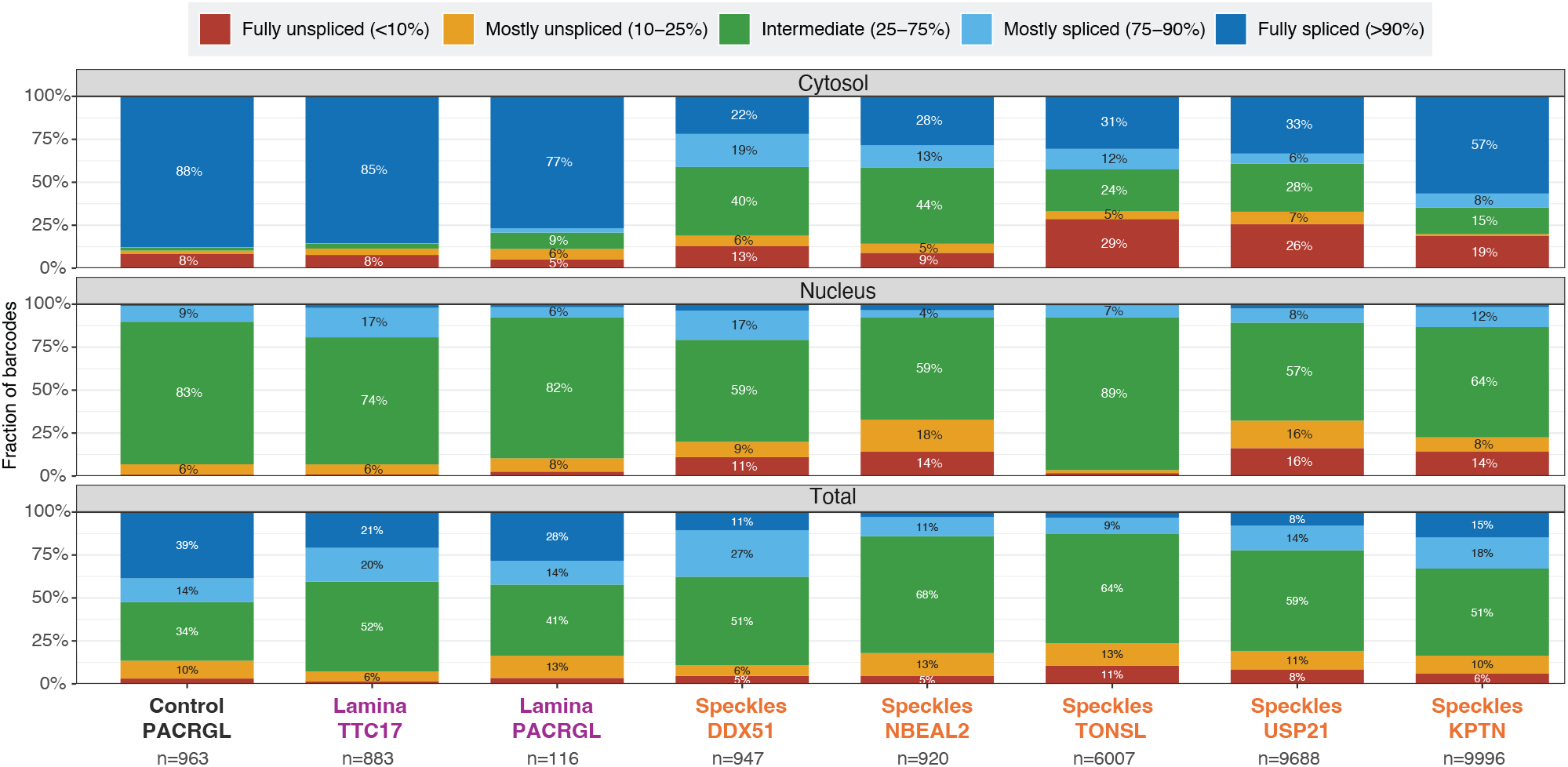
The localization fate of IntronTRIP reporters. Distribution of splicing efficiencies for barcodes of the indicated IntronTRIP reporter in the cytosol, nuclear, and total (whole cell extract) fractions. We considered only barcodes with at least 20 reads in each fraction, and we indicate the number of usable barcodes per reporter beneath.

## Discussion

Separating sequence-intrinsic effects from regulation imposed by the local genomic context is a central challenge in understanding gene regulation. In this study, we used TRIP to experimentally disentangle sequence-intrinsic properties of RNA processing from the genomic environments in which they are normally embedded. We find that genomic context strongly shapes transcriptional output and tunes RNA processing, whereas partially-retained introns preserve characteristic, portable behaviors across genomic locations.

Our approach allowed us to systematically dissect the relative contributions of genomic context and sequence on splicing. Although genomic position measurably tunes splicing outcomes, intron sequence remains the dominant determinant of splicing behavior, consistent with a previous study that tested different introns in a single genomic integration site (Mikl et al. 2019). Importantly, the magnitude of context dependence differs between introns, indicating that genomic features are not interpreted uniformly, but interact with intron-specific cis-regulatory features. Such sequence-dependent responses may arise through differential recruitment of RNA-binding proteins or through intron-specific interactions with the local chromatin environment. Both histone modifications and DNA methylation have been linked to intron retention and alternative splicing (Wong et al. 2017), and consistent with our observations, a previous study identified combinatorial and position-dependent chromatin signatures that are associated with alternative splicing, with similar chromatin features linked to different splicing outcomes depending in part on underlying RNA-binding motifs (Agirre et al. 2021). Together, our results support a model in which chromatin does not impose a universal effect on splicing, but instead tunes intron-intrinsic splicing patterns.

This sequence-dependent responsiveness to genomic context was particularly evident for nuclear speckle-associated introns. Nuclear speckles have emerged as important regulators of pre-mRNA processing, with speckle proximity associated with increased splicing efficiency (Bhat et al. 2024), and a distinct subset of partially retained introns depends on nuclear speckle-associated proteins for efficient splicing (J. Wu et al. 2024; Barutcu et al. 2022; Małszycki et al. 2026; Wen et al. 2026). Our results extend these observations by showing that speckle-associated introns retain characteristic processing behavior after redistribution across thousands of genomic positions. This suggests that reliance on SON, which is required for nuclear speckle formation, is not solely imposed by the host gene or genomic locations, but is at least partly encoded within this special class of introns.

Among the speckle-associated introns, DDX51 stands out as the one with the most distinguishing features, which manifest both at the level of its expression and splicing. Among the ten promoterless reporters in this study, it was the one with the broadest expression, with almost 60% of its barcodes being expressed (Fig. S4C), driven in large part by expression in weak enhancer state regions (Fig. S4B), while its expression was still low in repressed chromatin states. A plausible model is that the sequences of the exons and/or the intron in this reporter are more capable than others in engaging the RNA transcription and processing machinery and boosting gene expression, as part of the exon-mediated activation of transcription starts (EMATS) phenomenon (Uriostegui-Arcos et al. 2023). The splicing efficiency of the DDX51 intron also depends on features distinct from other introns in that, while it is more efficiently spliced near speckles, which are generally associated with GC-rich sequences and with high transcription, its splicing is relatively low in promoter- and enhancer-proximal states, and instead are highest away from transcription start sites, in typical weak/strong transcription states, associated with H3K36me3, and with low H3K4 methylation. Whether the elements that affect its broad expression are related to the elements that sculpt its splicing patterns is an interesting question for future studies. Interestingly, the DDX51 intron also exhibits cell-cycle-associated variation in intron retention in its endogenous context (Dominguez et al. 2016) and disruption of the important cell cycle kinase families CLKs and DYRKs leads to increased intron retention in DDX51 (E. Wang et al. 2023), consistent with the role of the DDX51 protein during cell cycle progression (X. Wang et al. 2016). Cell-cycle regulated introns are enriched among those showing increased retention upon SON knockdown (Barutcu et al. 2022). This may partly explain the stronger relations between speckle proximity and splicing observed for the DDX51 intron across genomic positions when compared with other introns.

We quantified intron retention in different cellular fractions and found that four of the five nuclear speckle-associated introns remained partially retained in host transcripts in the cytoplasm, mirroring their behavior in the endogenous transcript. The fifth speckle-associated intron, from KPTN, was instead more strongly restricted to the nucleus, both endogenously and across thousands of ectopic positions. Although intron retention often functions to keep transcripts in a poised state in the nucleus, awaiting signals for rapid splicing, export, and translation (Boutz et al. 2015), cellular surveillance mechanisms also act to degrade intron-retaining transcripts and restrict their abundance. In our panel, NBEAL2 and DDX51 are the only genes whose retained intron isoforms are selectively degraded by the nuclear exosome (Ietswaart et al. 2024). That these isoforms nonetheless reach the cytoplasm from thousands of ectopic positions indicates that nuclear surveillance does not fully sequester them once the intron is moved out of its native locus. Whether the degradation of the retained intron is itself intron-encoded, or depends on the native host gene, remains open.

Our approach has several limitations – at least 40% of the barcodes land in nontranscribed regions and therefore yield limited information; the integrations are biased toward more open chromatin regions; and every reporter is integrated into a different set of positions, making comparisons between different reporters challenging. We address the first limitation by integrating relatively large numbers of reporters compared to the original TRIP approach (Akhtar et al. 2013), which resulted in tens of thousands of expressed reporters. The two latter limitations partially address one another, in that the number of positions where different reporters are integrated is in the thousands. However, these shared positions are more strongly biased towards particularly active and accessible chromatin due to a ‘quadratic’ bias imposed by two separately biased integration events. The landing pad approach we devised partially resolves this problem, while the initial transposition of the landing pad is still biased by piggyBac preferences, subsequent recombination is much less biased, resulting in a distribution of chromatin states similar to that of individual reporter integrations (Fig. S6A). We initially integrated a relatively large number of landing pads, but limiting the number of integrations is possible and can allow for study designs in which dozens or more reporters can be integrated into shared sites.

Taken together, our results support a model in which RNA processing elements differ fundamentally in their autonomy. A cryptic polyadenylation site is defined by the region and is largely inert when moved, at least for the one site tested here. In contrast, specialized introns carry intrinsic information regarding their splicing, effect of surrounding chromatin context, and proximity to nuclear speckles. Short DNA elements can likewise act autonomously to position a locus at speckles through transcription factor binding sites (Chaturvedi et al. 2026). Thus, the recruitment of a locus to nuclear speckles and the speckle dependence of intron splicing appear to be encoded by short but distinct sequence elements within these introns.

## Methods

### Cell culture

K562 cells were maintained in Iscove’s Modified Dulbecco’s Medium (IMDM) and HeLa cells in Dulbecco’s Modified Eagle Medium (DMEM), both supplemented with 10% FBS and 1% penicillin/streptomycin. Cells were grown at 37 °C in a humidified 5% CO₂ incubator.

### TRIP plasmids construction

All plasmids were constructed by restriction digestion and ligation and assembled in pBluescript SK(+). Individual fragments were synthesized by Twist Bioscience, except for the 1 kb GNAI1 intronic fragment, which was synthesized by GenScript. All plasmids were verified by whole-plasmid sequencing (Plasmidsaurus). Plasmid sequences are listed in the supplementary information, and intronic sequences in Table S9. Barcoded plasmid pools were generated essentially as described (Akhtar et al. 2014). Briefly, the insert was amplified with a forward primer carrying an 18-nt degenerate sequence, and the resulting barcoded amplicon was cloned into the parental TRIP plasmid by restriction–ligation, destroying a BglII site. Residual non-barcoded plasmid was then linearized by BglII digestion for 4 h, and the plasmid was purified, concentrated, and transformed into E. cloni Classic electrocompetent cells (Lucigen, 60117-1). Colonies were scraped from plates in PBS and plasmid DNA was extracted using the ZymoPure II Plasmid Maxiprep kit. Presence of the degenerate barcode was confirmed by Sanger sequencing of plasmid DNA from individual colonies and from a small pool of colonies.

### Generation of standard TRIP pools

TRIP pools were generated according to the original protocol (Akhtar et al. 2014) with minor modifications. Briefly, 6 × 10⁶ cells were transfected with 5 µg of a plasmid encoding piggyBac transposase and mCherry, together with 30 µg of the TRIP library, using Lipofectamine 2000. Twenty-four hours post-transfection, mCherry-positive cells were isolated by fluorescence-activated cell sorting (FACS). Cells with very strong mCherry signal were discarded to minimize the number of cells carrying multiple integrations. Sorted cells were then incubated with 0.5 µM 4-hydroxytamoxifen (4-OHT) for 24 h and expanded for five days, after which two sub-pools of 20,000 cells each were subcultured to establish biological replicates; for the promoter TRIP pool, sub-pools of 25,000 cells were used. These pools were grown for a further seven days, split into two new sub-pools to serve as technical replicates, and grown for another seven days before genomic DNA and RNA were extracted. Integration sites were mapped as previously described (Akhtar et al. 2014). Genomic DNA was digested with DpnII, purified, and self-ligated. Ligated DNA was purified, and inverse PCR was used to amplify barcodes together with their flanking genomic sequence, with six independent PCRs per TRIP pool. Mapping was performed twice per pool to define a set of high-confidence integration positions. To validate these assignments, a randomly selected subset of sites was confirmed by two rounds of nested PCR.

### Landing pad (LP) TRIP

Landing pad TRIP pools were established essentially as described (Hong et al. 2024). The landing pad fragment and the TagBFP2 reporter gene were either synthesized by Twist Bioscience or amplified from existing plasmids, and assembled by restriction–ligation cloning. The resulting plasmid was barcoded and transfected into K562 cells as described above. After seven days, individual LP pools were established and expanded. Barcoded intron plasmids were then nucleofected into the LP pools together with plasmids expressing φC31–mNeonGreen and Bxb1–mScarlet-I. After 24 h, triple-positive (TagBFP2, mNeonGreen, mScarlet-I) cells were sorted by FACS and expanded for seven days before harvesting for DNA and RNA extraction. Genomic DNA was used to make normalization and mapping libraries as described for regular and intronTRIP. Furthermore, we prepared amplicon libraries that capture both the intron barcode and the LP barcode using the LP_mapping primer and the corresponding normalization primers, as listed in Table S8.

### SON degron TRIP

Endogenous SON was tagged with HaloTag using plasmids described previously (Małszycki et al. 2026): a construct containing the HaloTag flanked by homology arms and a construct expressing the gRNA and Cas9 were nucleofected into K562 cells at a 2:1 ratio using a Lonza 4D-Nucleofector and the SF Cell Line kit according to the manufacturer’s instructions. After 24 h, cells were placed under puromycin selection (1-2 µg/ml) for 72 h to remove non-nucleofected cells. Single cells were then plated and expanded for 14 days without selection. Homozygously tagged clones were identified by PCR genotyping, DNA sequencing (Plasmidsaurus), and RNA analysis. TRIP pools were established in a homozygous clone as described for standard TRIP. For depletion of HaloTagged SON, 4 × 10⁶ cells were treated with 1 µM HaloPROTAC3 (MedChemExpress, #HY-111997) or DMSO vehicle control for 16 h.

### Genomic DNA extraction

High-quality genomic DNA was extracted as previously described (S. Chen et al. 2015) with minor modifications. Briefly, 4 × 10⁶ cells were harvested and resuspended in 600 µl NK lysis buffer (50 mM Tris-HCl pH 8.0, 50 mM EDTA, 1% SDS). Proteinase K (3 µl of 20 mg/ml) was added and samples were incubated overnight at 55 °C. The following day, 3 µl RNase A (10 mg/ml) was added and samples were incubated for at least 30 min at 37 °C. Samples were cooled, 200 µl ice-cold 7.5 M ammonium acetate was added, and samples were vortexed vigorously for 20 s and centrifuged at 4,000 × g for 10 min. The supernatant was transferred to a fresh tube, mixed with 600 µl isopropanol by inversion (50 times), and centrifuged at 4,000 × g for 10 min to pellet the DNA. The pellet was washed once with 1 ml 70% ethanol, briefly air-dried, and resuspended in 200 µl TE buffer. DNA concentration was measured on a Qubit fluorometer and integrity was assessed by gel electrophoresis.

### RNA extraction

RNA was extracted using TRI Reagent according to the manufacturer’s protocol and treated with Baseline-ZERO DNase as recommended by the manufacturer. RNA was reverse transcribed using the qScript FLEX cDNA Synthesis kit according to the manufacturer’s protocol, using either gene-specific primers or a mixture of random hexamers and oligo(dT) primers.

### Poly(A) selection

Poly(A) RNA was enriched as previously described (Despic and Jaffrey 2023). In brief, 25 µg total RNA was diluted in 75 µl water and mixed with 50 µl Oligo(dT)₂₅ beads (NEB, S1419S) in 75 µl 2× mRNA binding buffer (40 mM Tris-HCl pH 7.5, 1 M LiCl, 2 mM EDTA, 0.1% Triton X-100). Samples were heated at 65 °C for 5 min, placed on ice for 3 min, and then incubated at room temperature with rotation for 10 min. Beads were washed twice with 150 µl mRNA wash buffer (20 mM Tris-HCl pH 7.5, 150 mM LiCl, 1 mM EDTA, 0.01% Triton X-100), resuspended in 75 µl mRNA elution buffer (10 mM Tris-HCl pH 7.5), heated at 75 °C for 2.5 min, and placed on ice for 2 min. Eluted RNA was subjected to a second round of purification. 75 µl 2× mRNA binding buffer was added, and the sample was incubated for 10 min at room temperature with rotation, washed once with 150 µl mRNA wash buffer, and resuspended in 30 µl water. Samples were heated at 75 °C for 3 min, and the supernatant was collected. Of the resulting mRNA, 100 ng was used in a 20 µl cDNA reaction as described above. To recover the poly(A)-minus fraction, the supernatant from the first binding step was retained, ethanol precipitated, and resuspended in water. 1 µg of Poly(A)-minus RNA was used for cDNA synthesis.

### Preparation of amplicon libraries

Expression amplicons were amplified from cDNA with Phusion polymerase (NEB) in two rounds of PCR as previously described (Akhtar et al. 2014), using 2 µl cDNA in the first round. Four parallel reactions were performed for each biological replicate of TRIP. Normalization amplicons were amplified in parallel under the same conditions from 100 ng genomic DNA, also in four reactions. All amplicon PCRs included three sets of controls; -RT samples, wild-type template and water. PCR products were pooled and purified using NucleoSpin Gel & PCR Clean-up kit. Libraries were quantified on a Qubit fluorometer, and fragment size distribution was assessed on an Agilent TapeStation. Libraries were sequenced on an Illumina NovaSeq 6000 or NovaSeq X instrument to a depth of approximately 15 million reads per library.

### cDNA Mapping

Mapping of the RNA sequences flanking TRIP reporters was performed by adapting the TIF-seq2 protocol described in (B. Li et al. 2021). Total RNA was extracted using TRI Reagent and treated with Baseline-Zero DNase to remove residual genomic DNA. For upstream sequence mapping, RNA was dephosphorylated with alkaline phosphatase and decapped with Cap-Clip acid pyrophosphatase. After phenol:chloroform purification, the decapped RNA was ligated to a DNA/RNA chimeric adapter using T4 RNA ligase for 16 h at 16 °C. Reverse transcription was performed using SuperScript III reverse transcriptase in the presence of 1.57 M trehalose and a TRIP-specific primer. RNA templates were then removed by treatment with RNase H, RNase A and RNase T1 for 30 min at 37 °C. cDNA was purified using AMPure XP beads and converted to double-stranded cDNA using Terra PCR Direct polymerase and a TRIP-specific reverse primer containing an MspI recognition site. ds-cDNA was purified with AMPure XP beads and digested with MspI for 4 h at 37 °C. Fragmented DNA was purified using AMPure XP beads, diluted to 400 µl and ligated overnight at 16 °C to favor intramolecular ligation. Ligase was inactivated at 65 °C for 10 min, after which circularized cDNA was purified by phenol:chloroform:isoamyl alcohol extraction followed by ethanol precipitation. Circularized cDNA was used as the template for three rounds of inverse PCR, as described for DNA mapping. Mapping of downstream sequences was performed as above, with the following modifications: the dephosphorylation, decapping and adapter ligation steps were omitted; reverse transcription was primed with an oligo(dT) primer carrying a 5′ adapter sequence for subsequent PCR amplification; second-strand synthesis was done with a TRIP-specific forward primer; and ds-cDNA was digested with DpnII.

### Morpholino delivery

K562 and HeLa cells (1.5 × 10⁶ per reaction) were electroporated with 7.5 nmol (75 µM final) of an antisense morpholino oligonucleotide (AMO) targeting U1 snRNA (GGTATCTCCCCTGCCAGGTAAGTAT), or a scrambled control AMO (CCTCTTACCTCAGTTACAATTTATA), in a transfection volume of 100 µl using a Lonza 4D-Nucleofector with the SF and SE Cell Line kits, respectively. Electroporated cells were plated in 2 ml prewarmed medium and harvested 8 h after electroporation. U1 snRNP inhibition was confirmed by semi-quantitative RT-PCR of the U1-sensitive RAN transcript (Berg et al. 2012).

### NMD inhibition

2 million cells from the indicated TRIP pools were treated with 100 µg/ml cycloheximide (CHX; C7698, Sigma) or an equivalent volume of the vehicle control (ethanol) for 4 h. Following treatment, RNA was extracted, and expression amplicons were generated and sequenced as described above.

### RNA sequencing

Total RNA was extracted from U1 or scrambled AMO-treated cells as described above. PolyA RNA was enriched using the Poly(A) RNA Selection Kit V1.5 (Lexogen) according to the manufacturer’s instructions, and libraries were prepared using the CORALL mRNA-Seq V2 Kit (Lexogen) following the manufacturer’s recommendations. Total RNA was extracted from HaloPROTAC3- or DMSO-treated cells as described above. Poly(A) selected RNA was isolated using the NEBNext Poly(A) mRNA Magnetic Isolation Module (NEB) according to the manufacturer’s instructions, and libraries were prepared using the NEBNext Ultra II Directional RNA Library Prep Kit for Illumina (NEB) following the manufacturer’s recommendations. Libraries were sequenced on an Illumina NovaSeq 6000 or NovaSeq X.

### RT-PCR and RT-qPCR

RNA was purified as described above. Equal amounts of RNA were reverse transcribed using the qScript FLEX cDNA Synthesis Kit (Quantabio). cDNA was used as input for PCR with Phusion High-Fidelity DNA polymerase (NEB) according to the manufacturer’s specifications. For RT-qPCR, cDNA was diluted as appropriate and used as input for quantitative real-time PCR with Fast SYBR Green Master Mix (Thermo Fisher, 4385614) on a ViiA 7 Real-Time PCR System (Applied Biosystems).

### Subcellular fractionation

Fractionations were performed as previously described (Lubelsky and Ulitsky 2018), with minor modifications for K562 cells. Cells were collected, washed once with PBS, lysed in RLN buffer (50 mM Tris-Cl pH 8, 140 mM NaCl, 1.5 mM MgCl₂, 0.15% v/v NP-40, 10 mM EDTA, 1 mM DTT, 10 U/ml RNase inhibitor) and incubated on ice for 5 min. Fractions were separated by centrifugation for 5 min at 300 × g and 4 °C. The supernatant was collected and centrifuged for 1 min at 500 × g; the resulting cytoplasmic fraction was collected, and RNA was extracted with TRI Reagent. The nuclear pellet from the first centrifugation was washed once in S1 buffer (50 mM sucrose, 10 mM MgCl₂, 10 U/ml RNase inhibitor), layered on top of S3 buffer (880 mM sucrose, 0.5 mM MgCl₂, 10 U/ml RNase inhibitor) and centrifuged for 10 min at 2,800 × g and 4 °C. The supernatant was discarded, and nuclear RNA was extracted from the pellet using TRI Reagent. Successful fractionation was confirmed by measuring the relative distribution of MALAT1 (nuclear) and ACTB (cytoplasmic) by RT-qPCR as described above.

### Immunoblotting

Western blotting was performed as previously described (Roseman et al. 2025). Cells (1 × 10⁶) were harvested, washed once with ice-cold PBS and lysed on ice for 30 min in 50 µl RIPA buffer (50 mM Tris-HCl, 150 mM NaCl, 0.5% sodium deoxycholate, 0.1% SDS, 1% NP-40, 2 mM EDTA) freshly supplemented with EDTA-free protease inhibitor cocktail (APExBIO). Benzonase wash buffer (50 µl; 50 mM Tris-HCl, 50 mM NaCl, 2 mM MgCl₂, 1:500 Benzonase) was added, and samples were incubated for 20 min with rotation at room temperature. Lysates were clarified by centrifugation at 14,000 × g for 15 min at 4 °C. The supernatant was mixed with a loading buffer and heated for 10 min at 70 °C. Equal amounts of protein were resolved on a 6% polyacrylamide gel for 2 h. Transfer was performed on ice at a constant current of 0.30 A for 2 h in Tris-glycine transfer buffer containing 20% methanol, onto a pre-activated 0.45 µm PVDF membrane. After blocking, the membrane was incubated overnight at 4 °C with the primary antibody (Anti-SON 1:1000, HPA023535, Sigma) diluted in 5% milk in 1× PBS with 0.1% Tween 20. The next day, membranes were washed three times with 1X PBS-0.1% Tween 20, incubated for 1 hr at room temperature with the appropriate secondary antibody, and washed three more times before acquisition. For direct visualization of Halo-SON, HaloTag TMR ligand (Promega G8252) was added to a final concentration of 0.5 µM during protein extraction and the gel was imaged on a ChemiDoc prior to transfer.

### Read processing and integration-site mapping

All mapping, expression, and normalization libraries were trimmed with cutadapt. Barcodes were taken as the 18 bases lying between the two constant reporter sequences that flank them. Adapter sequences for every library type, including the isoform-specific adapters used to distinguish spliced from unspliced amplicons, are listed in Table S8.

#### Processing mapping files

We first removed the adaptors from the reads using cutadapt --cut 4 -g AGAAGTTCAACTCGGTACCAGAGCCACCNNNNNNNNNNNNNNNNNNGGATC --rename=’{id} umi={cut_prefix} adapter={adapter_name} barcode={match_sequence}’ -a ADAPTOR1 -o ./01_cutadapt/${p}.R1.fastq.gz -p ./01_cutadapt/${p}.R2.fastq.gz-G ADAPTOR2

Extracted the barcodes from each read by examining the 18 bases between AGAAGTTCAACTCGGTACCAGAGCCACC and `GGATC`, and then used Bowtie2 to map the remaining sequences with `-X 2000` parameter, converted the BAM files to BED files, using BEDtools bamtobed and extracted the beginning of the alignment of read2 as the integration site, i.e. the junction between the piggyBac ITR and the genome. For sites reported on the plus strand, the TTAA occupies the four bases immediately upstream (offset −4), and for minus-strand sites the four bases begin at the site itself (offset 0). Across the mapping libraries, >80% of retained barcodes fell at one of these two canonical positions (versus <1% for position-shuffled controls). Sites without a TTAA at the expected offset were predominantly barcodes supported by a single mapping read and were discarded.

#### Mapping barcode positions

Reads sharing a barcode were grouped by alignment position, and barcodes differing by up to three substitutions were merged into the more abundant variant. A candidate position was then discarded when it lay within 10 bp of a position supported by more reads, carried at most 25% of that position’s read count, and its top barcode was within edit distance 3 of the stronger position’s top barcode. As a barcode observed in more than one pool could indicate a potential cross-contamination during sequencing, for every barcode–position pair we compared its count in each pool against the highest count that the same barcode reaches at the same position in any other pool sequenced in the same sequencing run, and discarded the observation when it fell below that maximum by more than a factor of 10. Barcode–position calls were then pooled across the mapping replicates of each pool. Only barcodes of the expected length (18 nt; 16 nt for the landing-pad libraries). Within each position, read counts were summed per unique barcode and barcodes within a Levenshtein distance of 2 of a more abundant barcode were merged into it, absorbing residual sequencing error. A barcode mapping to exactly one genomic position was used for all downstream analysis; barcodes appearing at more than one position were set aside as multi-mappers and excluded from further consideration.

#### Annotation of barcode positions

The hg38 assembly was used throughout, with the exception of nuclear speckle distances, which were based on hg19. Integration sites were intersected strand-specifically with MANE Select v1.0 transcripts and their introns, with GENCODE v46 non-coding genes, and with RepeatMasker annotations, recording both sense and antisense overlaps. For each site falling within a MANE transcript we computed the distance to the TSS and to the TES, the relative position along the transcript, the distances to the 5′ and 3′ ends of the host intron, the relative position within that intron, and the host intron length. G/C content was computed over the 500 bp flanking each side of the integration site. Histone-mark ChIP-seq and additional genomic datasets were obtained as bigWig files from the ENCODE project.

#### Measurements of expression levels

Barcodes were counted from trimmed read1 of each expression and normalization library by extracting the 18-nt barcode at a fixed offset determined by the adapter used and tallying unique sequences RNA. RNA and DNA normalization barcode counts were matched per pool. A barcode was required to reach at least 2 reads in the DNA normalization library to be carried forward. Barcode expression was defined as log10(1 + reads), and when comparing libraries of different depth, the read count in each library was scaled to its own total depth.

#### Measurements of splicing efficiency

For each barcode, spliced and unspliced amplicons were counted separately using isoform-specific read 2 adapters (Table S8), and splicing efficiency was defined as spliced / (spliced + unspliced) reads, and was computed only for barcodes with at least 20 combined (spliced + unspliced) reads.

#### Processing of landing-pad libraries

Landing-pad mapping libraries were trimmed with a 5′ anchored adapter capturing the 16-nt genomic barcode on read 1 and the piggyBac ITR adapter on read 2; normalization libraries were trimmed with the corresponding anchored adapters for the landing-pad construct, and recombination libraries with reporter-specific anchored adapters, all listed in Table S8. In recombination amplicons, read 1 carries the 16-bp landing-pad barcode identifying the genomic position and read 2 carries the 18-bp reporter barcode, allowing each recombined reporter to be assigned to its mapped landing-pad site.

### Machine learning

Data preparation, model training and SHAP computation were performed in Python 3.11.13 with xgboost 3.0.3, shap 0.48.0, scikit-learn 1.7.1, numpy 2.2.6, pandas 2.3.1, scipy 1.16.0, intervaltree 3.0.2, matplotlib 3.10.0 and seaborn 0.13.2. Code for model training, group SHAP and the NMD feature is available at https://github.com/ArtyomTyukaev/trip-ml-part. The full software environment specification is included in the repository.

## Supporting information

Plasmids

## Acknowledgements

We thank Noam Stern-Ginossar, Schraga Schwartz, Dana Segal, and members of the Ulitsky lab for helpful discussions and comments on the manuscript.

## Data availability

All sequencing libraries were deposited in GEO under accessions GSE352314 (regular TRIP; reviewer access token obulgoaapvopjsf); GSE352315 (IntronTRIP, reviewer access token srefqaoinriltov); GSE352316 (Landing pad TRIP experiments; reviewer token slqxsassznuvhcv); GSE352317 (U1 inhibition experiments; reviewer access token svarwiwadfubtsx); GSE352318 (CHX treatment of TRIP libraries, reviewer access token ktyxqqkitjivfir); GSE352319 (SON depletion, ubyzmocgtbahhsx); GSE352321 (CHX treatment of IntronTRIP libraries, reviewer access token kronwgugrrwtfup). The full list of the sequencing libraries is found in Table S10.

## Code availability

The pipelines for data processing used in this study are available here: https://github.com/IgorUlitsky/SpliceTRIP; machine learning pipelines are available at https://github.com/ArtyomTyukaev/trip-ml-part.

## Supplementary Figures

**Figure S1.**
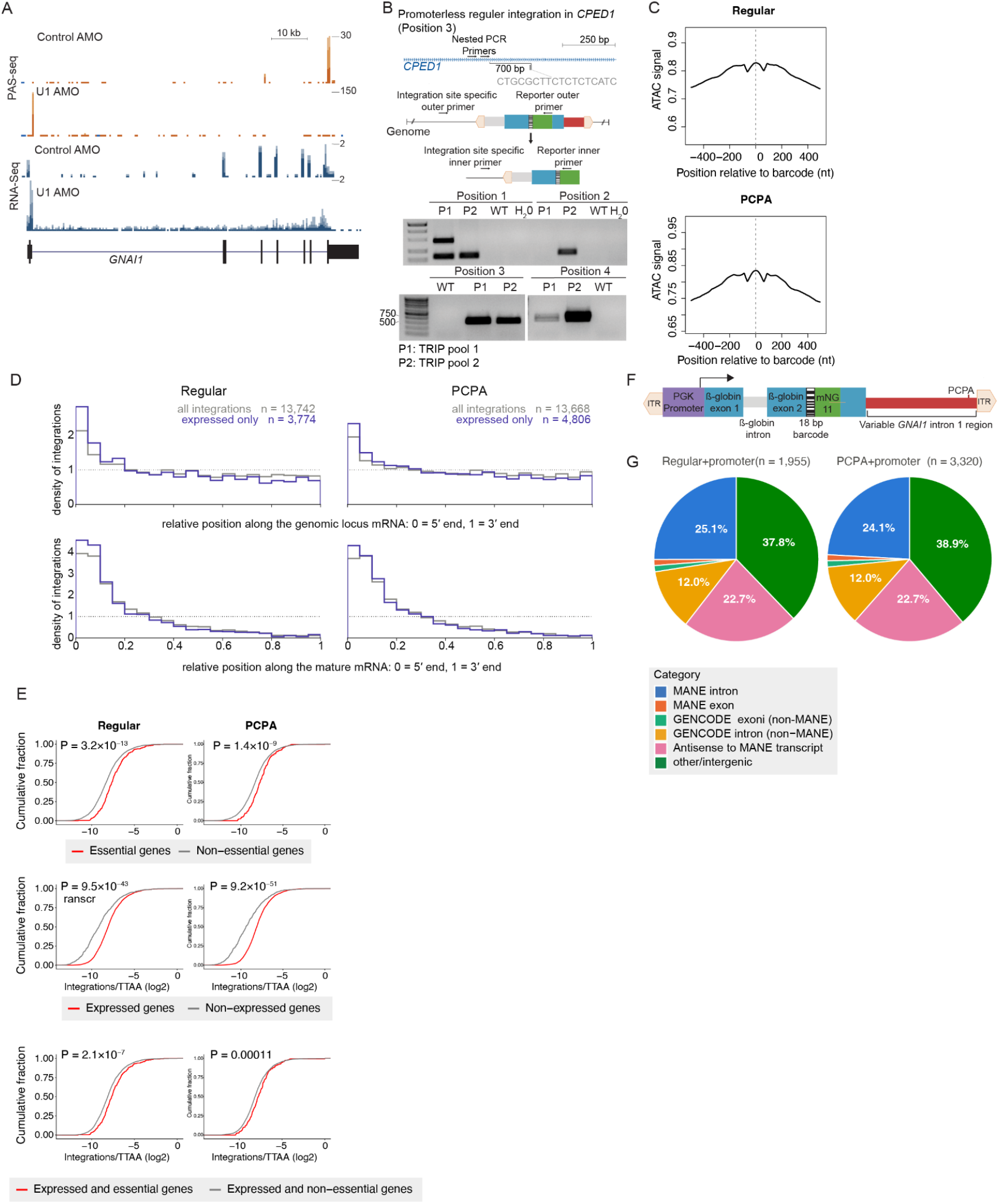
(A) PolyA site (PAS) sequencing and RNA-seq in HeLa cells at the *GNAI1* gene, with the early PCPA site used in the PCPA reporters. **(B)** Validation of genomic positions for four selected positions. Top: illustration of the PCR strategy; Bottom: PCR products obtained in the indicated pools. **(C)** Metagene of the ENCODE K562 ATAC-seq read coverage around the barcode integration positions for the Regular and PCPA reporters. **(D)** Density of integrations in the indicated bins across the MANE gene bodies (top) and MANE mRNAs (bottom). **(E)** Comparison of the density of barcode integrations as a function of all TTAA genomic sites in the indicated groups of genes. Essential genes are those with essentiality scores lower than –1 in K562 in DepMap. **(F)** Illustration of the structure of the promoter-containing Regular and PCPA reporters. **(G)** Fraction of barcodes in each library mapping to the indicated genomic region.

**Figure S2.**
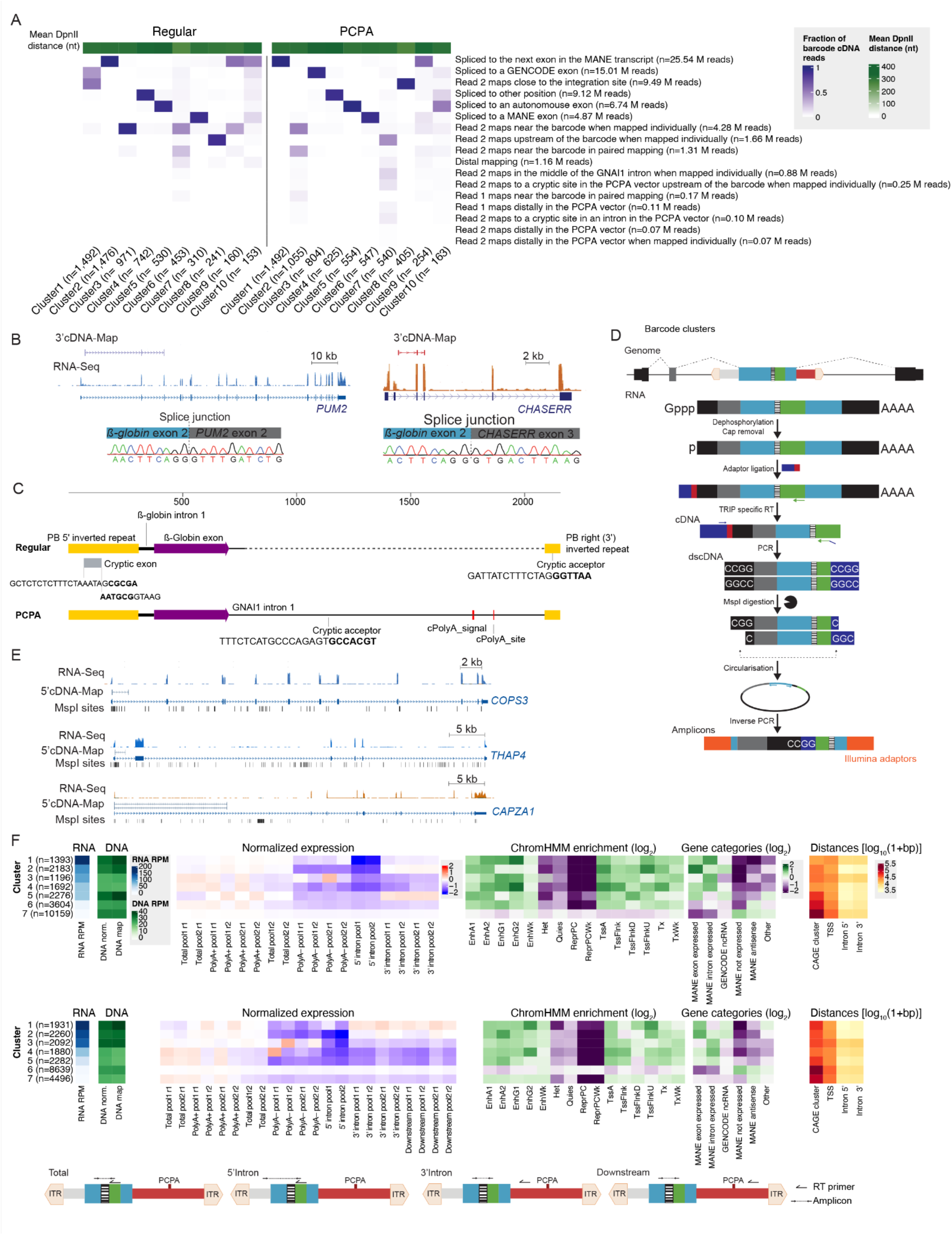
Additional information on integration of reporter exons into host genes. **(A)** Overview of the patterns of reads for reporter-host 3’ mapping. **(B)** Validation of the formation of splice junctions and host exons in the indicated genes with Sanger sequencing of the PCR product formed across the splice junction shown. (C) Outline of the reporter structure, with elements drawn to scale. The cryptic exon observed in the 5’ inverted repeat and the cryptic 3’ splice sites are shown, alongside the annotated PCPA site in the GNAI1 intron. Sequences of select splice junctions are shown with exonic sequence in bold. **(D)** Overview of the approach for mapping reporter-host 5’ connectivity. **(E)** Validation with 5’ reporter-host connectivity mapping at three integrations. **(F)** Expression-pattern clusters of TRIP reporter integrations and their genomic context. Barcodes were grouped by k-means clustering with k=7 for their expression profile across libraries. We clustered barcodes quantified in every library of the set (no missing values) and detected (non-zero) in at least 10% of the total expression libraries. Per library, counts were depth-normalized to reads-per-million (RPM); each barcode’s profile was then centred by subtracting its mean across the total expression libraries, so values represent expression relative to that barcode’s own average expression level. The panels show (left to right): i) Average RPM per cluster across the total RNA libraries, before the per-barcode centring used for clustering; ii) Mean RPM-normalized DNA reads in the normalization and map libraries; iii) k-means cluster centres, each column shows log ratio of that library to the barcode’s mean across the total expression libraries; iv) Enrichment of each ChromHMM chromatin state with ≥50 barcodes within each cluster, as log₂ of observed over expected, where expected is based on the state’s genome-wide frequency among all annotated barcodes; (v) Same for gene annotation classes; “expressed” refers to host genes above a normalized-expression threshold; (vi) Distances of the barcode to the indicated landmark (only for barcodes where the relevant landmark is available, e.g., distances to intron edges are computed only for barcodes within MANE introns in a sense orientation. Per cluster, the mean of log₁₀(1+distance in bp) for: distance to the closest upstream/overlapping FANTOM5 K562 CAGE peak (strand-specific), and distance to the TSS, to the 5′ end of the host intron, and to its 3′ end. Distances are log-transformed before averaging so that a few very distant barcodes do not dominate. Darker = further away. The schematic below the plot shows the combination of RT primer and PCR primers

**Figure S3.**
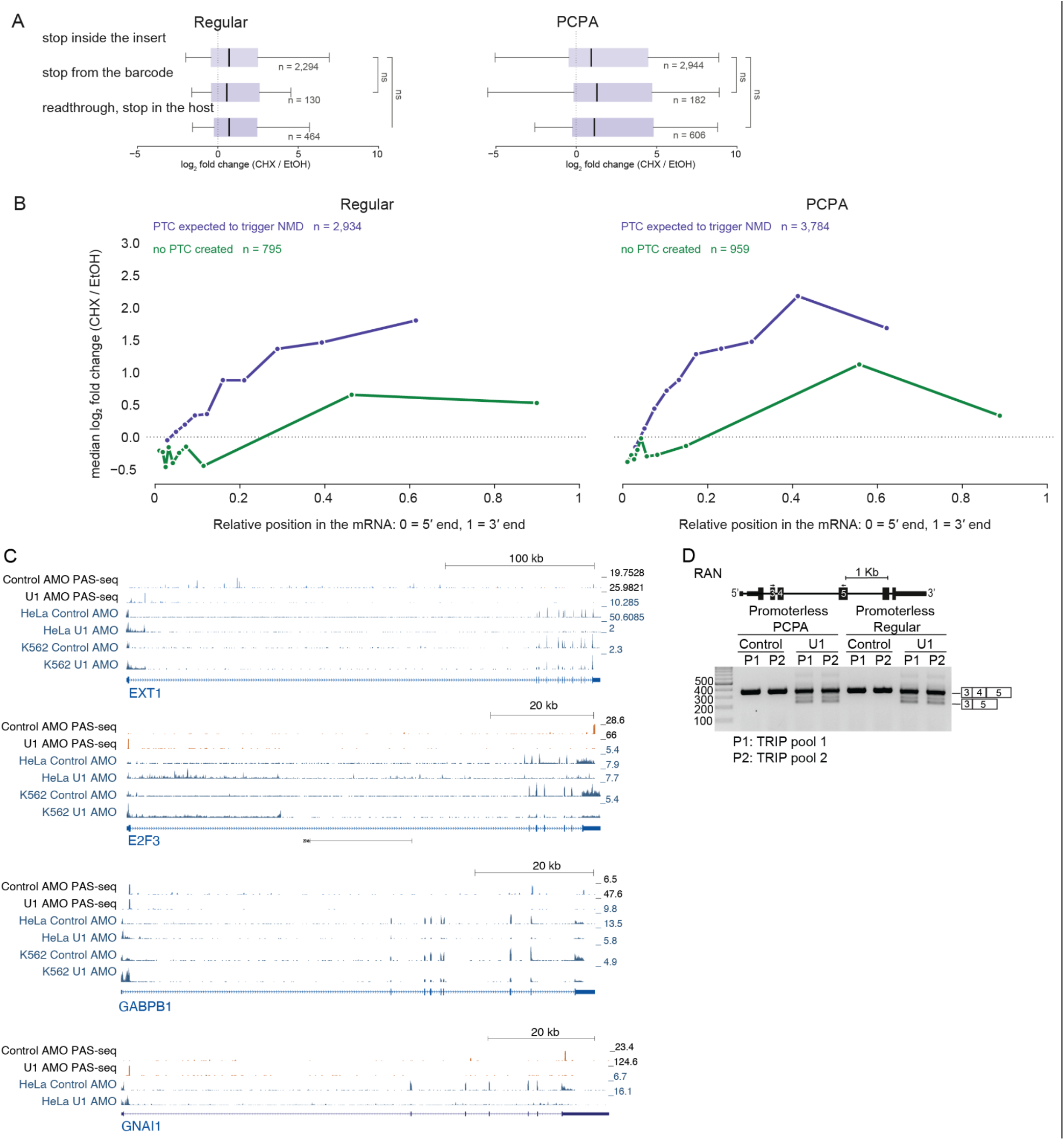
Effects of CHX and U1 AMO on the transcriptome. **(A)** Changes in gene expression of barcodes in the indicated library upon treatment of cells with CHX or the EtOH vehicle control for 4 hr, grouping the barcodes based on the nature of the PTC that is predicted to lead to NMD. n.s. stands for P>0.05 when using Wilcoxon rank-sum test. **(B)** CHX-induced change in barcode expression versus relative integration position along the mature host mRNA (0 = 5′, 1 = 3′). Integrations were grouped into 10 equally populated bins. Points show bin medians. PTC/NMD-triggering integrations are shown separately from those not creating a PTC. **(C)** Comparison of PAS-seq and RNA-seq read coverage following the introduction of control AMO or U1-targeting AMO in HeLa cells (PAS-seq (Shi et al. 2019) and RNA-seq tracks) and K562 cells (RNA-seq). The RNA-seq data are from this study (see Methods). MANE gene models are shown. **(D)** RT-PCR of the mis-splicing event in the RAN transcript following U1 AMO treatment(Berg et al. 2012).

**Figure S4.**
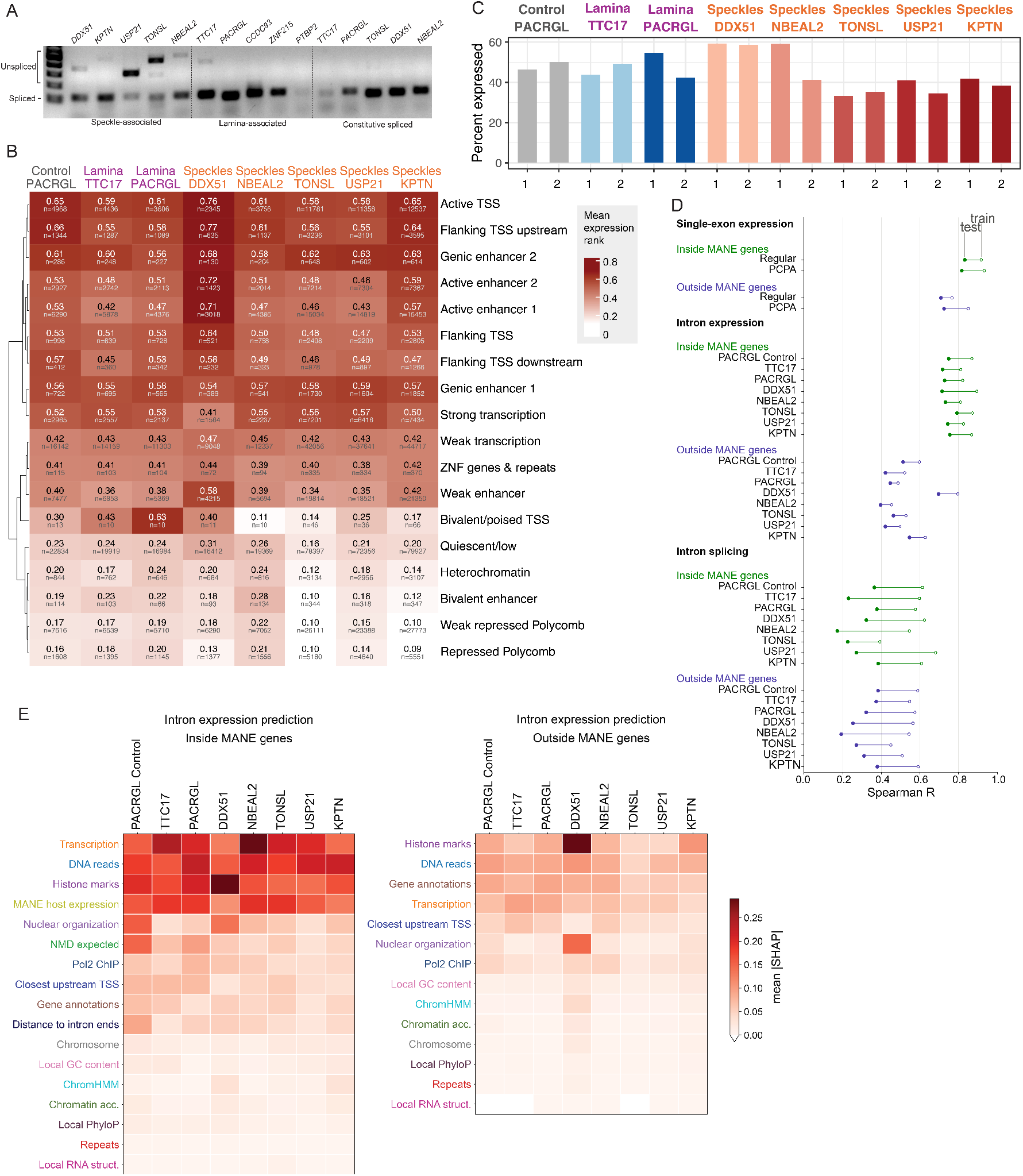
Information about the expression of IntronTRIP libraries. **(A)** RT-PCR on the indicated introns from the three groups. Primers are listed in Table S8. **(B)** Average expression rank for IntronTRIP barcodes split by ChromHMM state. For barcodes present in both pools, the average is shown (1=most reads; 0=least reads; for inserts with equal rank, the lowest rank is given). **(C)** Fraction of barcodes for every intron and pool detected with at least one read in the total expression libraries. **(D)** Spearman correlation coefficients between predicted and observed values for expression and splicing in the indicated TRIP or IntronTRIP assay. **(E)** SHAP values for the indicated feature sets in XGBoost predictions of total expression of the indicated intron, separately for integrations inside and outside MANE gene boundaries.

**Figure S5.**
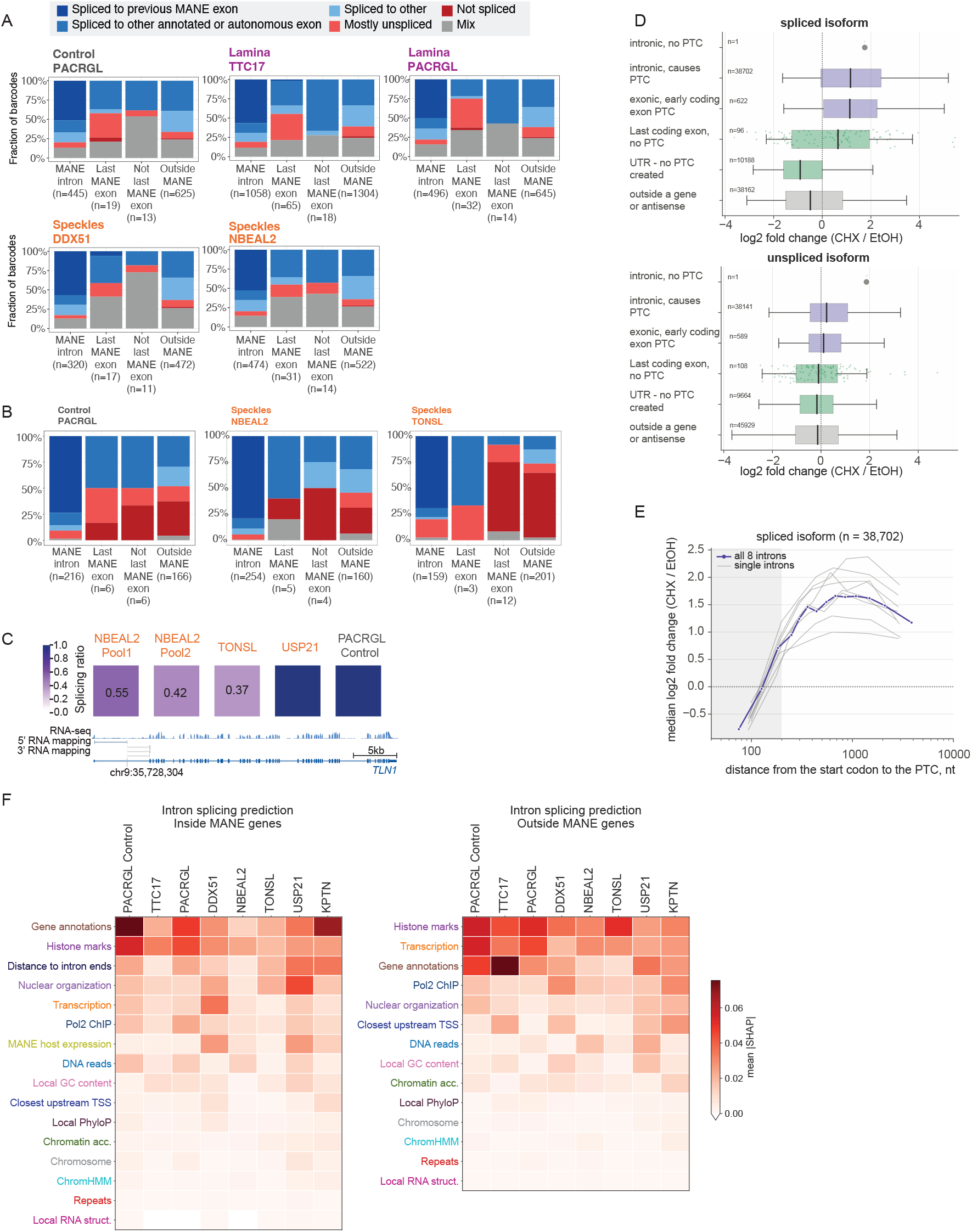
Additional information on the splicing of IntronTRIP libraries. **(A)** Distribution of reporter-host 3’ connectivity patterns observed for individual barcodes for which at least five informative reads could be obtained for each of the five exons for which such was performed. **(B)** Distribution of reporter-host 5’ connectivity patterns observed for individual barcodes for which at least five informative reads could be obtained for each of the five exons for which such was performed. **(C)** Example of a barcode position with both 5’ and 3’ connectivity support. Different barcodes were independently mapped in this position for each of four introns. The splicing efficiency for the indicated introns are shown. (**D)** Changes in expression of barcodes in the indicated isoforms upon treatment of cells with CHX or the EtOH vehicle control for 4 hr, grouping the barcodes based on position and the nature of the PTC that is predicted to lead to NMD. **(E)** CHX-induced change in spliced barcode expression versus distance from that start codon to the PTC. **(F)** SHAP values for the indicated feature sets in XGBoost predictions of splicing efficiency of the indicated intron, separately for integrations inside and outside MANE gene boundaries.

**Figure S6.**
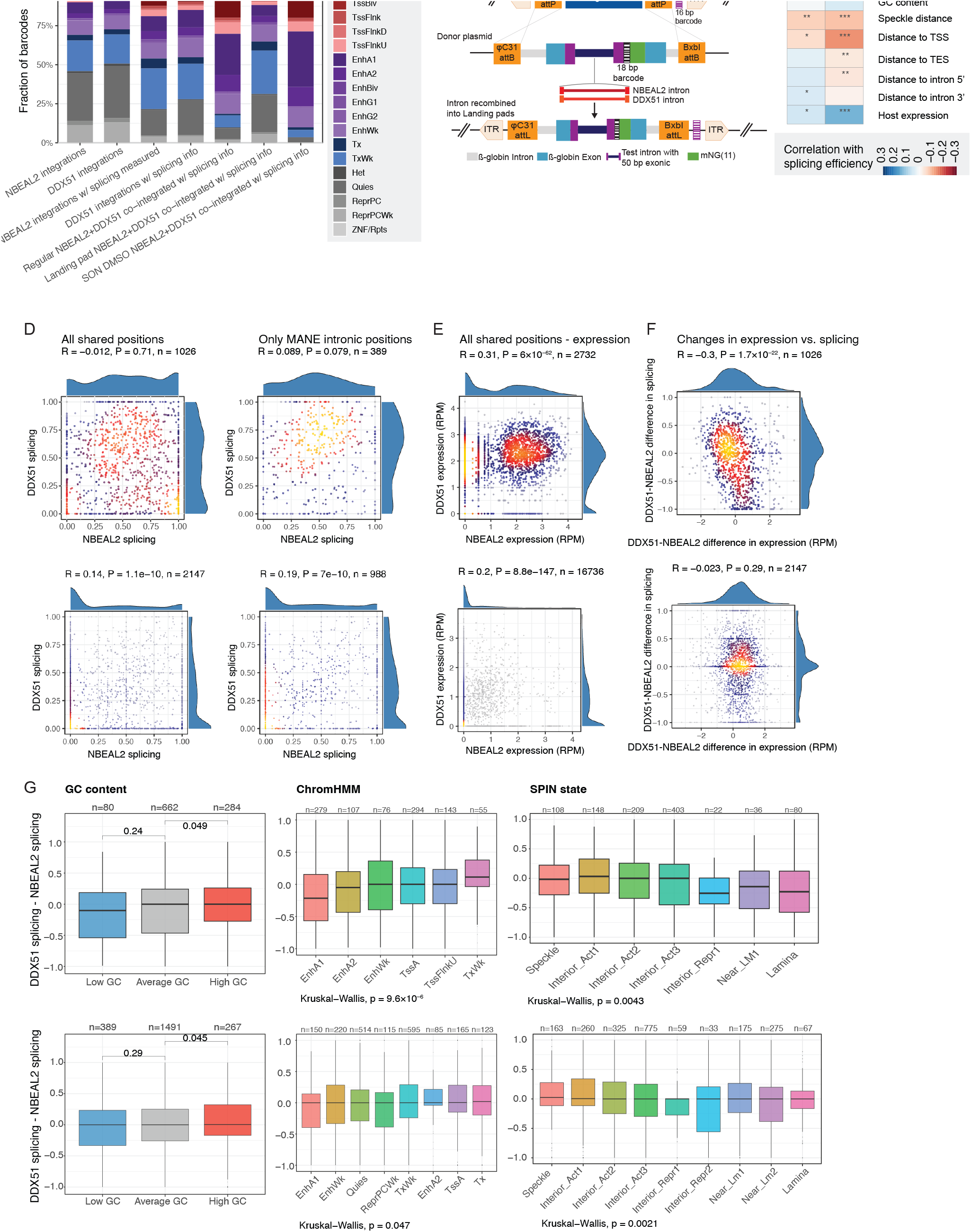
Correspondence between barcodes co-integrated into the same positions. **(A)** Distribution of ChromHMM states for the indicated set of integrations. **(B)** Outline of the approach for landing pad-based TRIP assay. **(C)** As in Fig. 5C, for the NBEAL2 and DDX51 introns integrated into the landing pad. **(D)** Correspondence of splicing efficiencies when using regular IntronTRIP (top) or landing pad IntronTRIP (bottom) at either all shared positions or just positions mapped inside MANE genes. **(E)** As in D for expression levels. **(F)** Correspondence between differences in splicing and expression levels between the two introns using regular IntronTRIP (top) or the landing-pad approach (bottom). **(G)** Distributions of differences in splicing between the two introns. P-values for G/C content classes are computed using the Wilcoxon rank-sum test. P-values for the effect of ChromHMM and SPIN states are computed using Kruskal-Wallis test.

**Figure S7.**
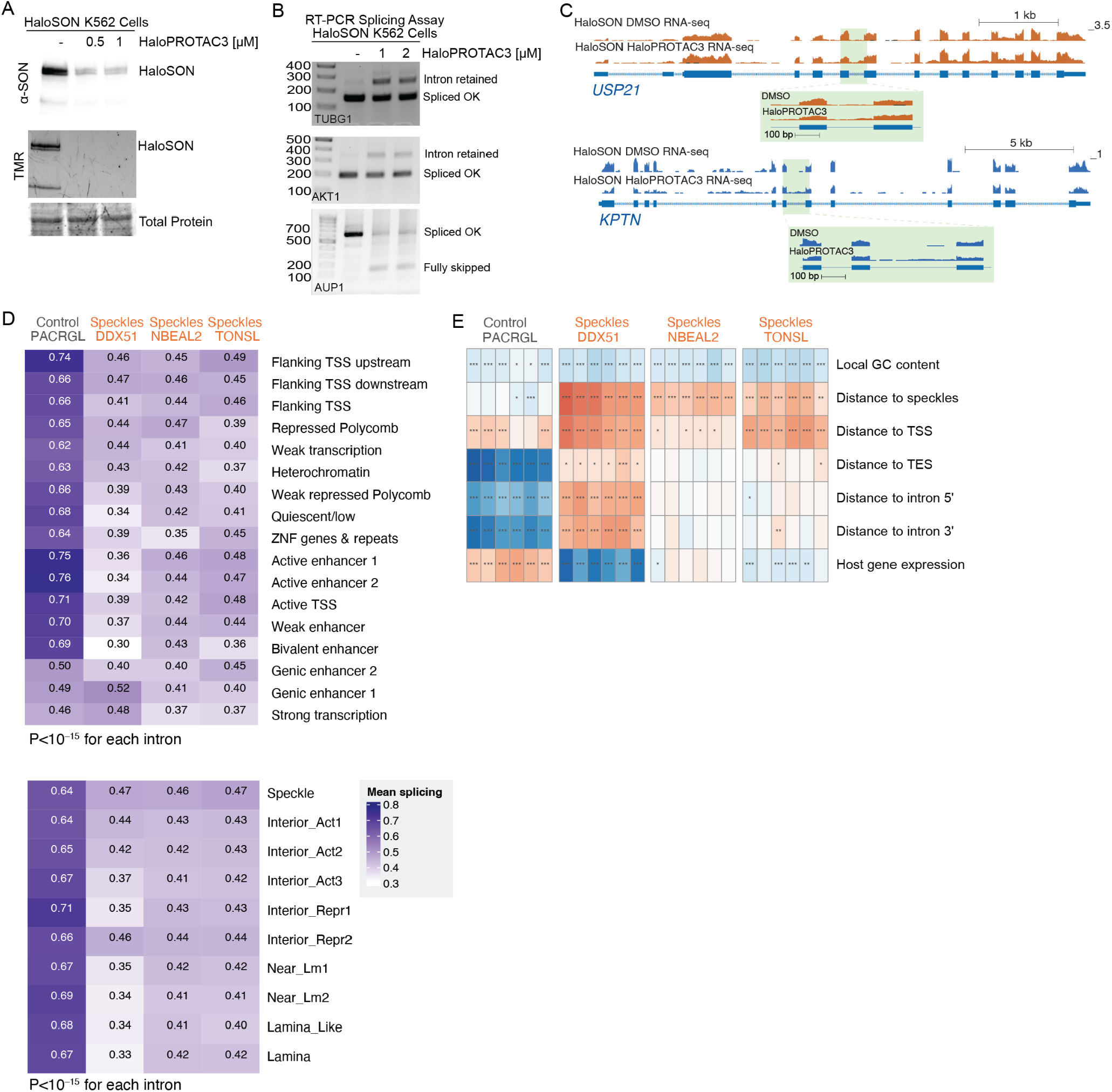
Characterization of HaloSON K562 cells. **(A)** Depletion of SON protein upon treatment with HaloPROTAC3 reagent. **(B)** Effects of SON depletion on the splicing of specific introns. **(C)** RNA-seq coverage on the endogenous USP21 and KPTN genes from which reporter introns were taken upon SON depletion in K562 cells. Zoom-in on the TRIP introns are shown below. **(D)** Mean splicing efficiencies of the indicated introns in the HaloSON K562 cells. **(E)** Correlations between splicing efficiencies and the indicated integration features.

**Figure S8.**
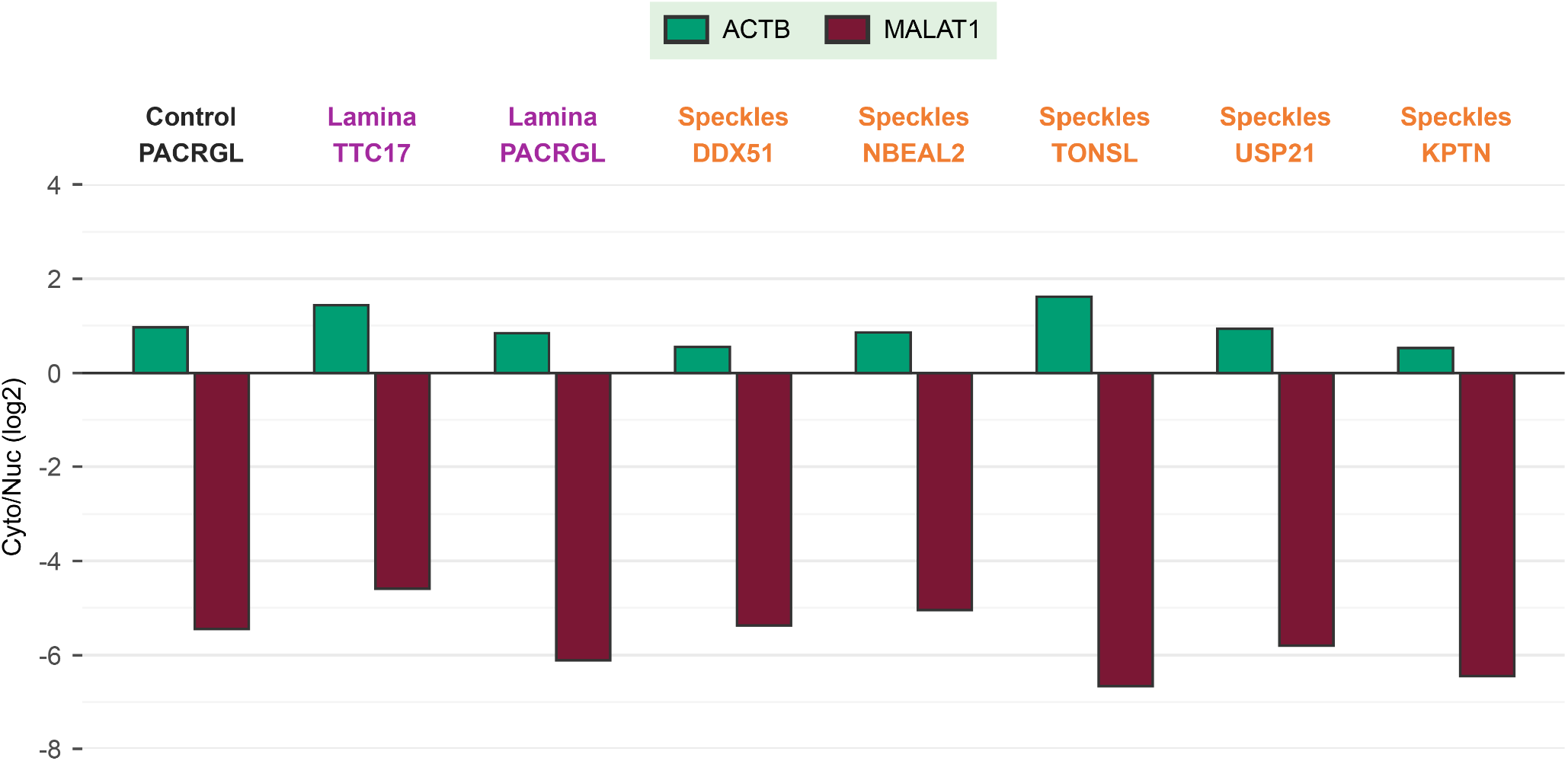
Validation of successful fractionation of IntronTRIP pools. **(A)** qRT-PCR for the Actin (ACTB) and MALAT1 in the indicated TRIP pools, comparing the cytoplasmic and nuclear enrichment.

## Supplementary Tables

Table S1. Barcode information for the Regular library

Table S2. Barcode information for the PCPA library

Table S3. Barcode information for the promoter libraries

Table S4. Barcode information for the IntronTRIP libraries

Table S5. Barcode information for the CHX-treatment IntronTRIP libraries

Table S6. Barcode information for the landing pad IntronTRIP libraries

Table S7. Barcode information for the HaloSON IntronTRIP libraries

Table S8. Primers and oligonucleotides

Table S9. List of intron sequences for IntronTRIP

Table S10. Summary of the sequencing libraries

